# Single cell whole genome and transcriptome sequencing links somatic mutations to cell identity and ancestry

**DOI:** 10.1101/2025.10.28.685157

**Authors:** Abhiram Natu, Mrunal K. Dehankar, Reenal Pattni, Milovan Suvakov, Dmitrii Olisov, Livia Tomasini, Yeongjun Jang, Yiling Huang, Eva Benito-Garragori, Patrick Hasenfeld, Jan O. Korbel, Alexander E Urban, Alexej Abyzov, Flora M. Vaccarino

**Affiliations:** Child Study Center, Yale University, New Haven, CT, 06520, USA; Program in Neurodevelopment and Regeneration, Yale University, New Haven, CT, 06520, USA; Department of Quantitative Health Sciences, Center for Individualized Medicine, Mayo Clinic, Rochester, MN, 55905, USA; Bioinformatics and Computational Biology Program, University of Minnesota, Minneapolis, MN 55455, USA; Department of Psychiatry and Behavioral Sciences, Stanford University, Palo Alto, CA 94305, USA; Department of Genetics, Stanford University, Palo Alto, CA 94305, USA; European Molecular Biology Laboratory (EMBL), Genome Biology Unit, 69117 Heidelberg, Germany; Department of Neuroscience, Yale University, New Haven, CT, 06520, USA; Kavli Institute for Neuroscience, Yale University, New Haven, CT, 06520, USA; Yale Stem Cell Center, Yale University, New Haven, CT, 06520, USA

## Abstract

The role of somatic mutations in human development and disease is obscured by difficulties in characterizing mutations at the single cell level and identifying cell types carrying them. Here we analysed somatic genomes of clonal iPSC lines and of single-cells after whole-genome amplification (scWGA) by PTA and ResolveOme from skin fibroblasts, blood and urine of a live donor. Mutation burden and spectra converged across approaches, revealing heterogeneous mutational footprints across cells driven by environmental exposures (UV damage and chemotherapy) and lymphocyte differentiation. Aneuploidies in single cells were detected by all the approaches and were orthogonally validated by Strand-seq. Uniquely, ResolveOme enabled cell-type identification using single-cell transcriptomes. Using a newly developed method accounting for noise and allele drop-out in scWGA, we *de novo* reconstructed the cell phylogenetic tree for this donor. Together, scWGA establishes a powerful foundation for comprehensive, cell type-aware, lineage-aware profiling of somatic mutations at single cell level.

## Introduction

Somatic mosaicism has emerged as a fundamental feature of human biology. Recent studies revealed that virtually every cell in the human body harbours a unique constellation of somatic mutations, ranging from single-nucleotide variants to complex structural alterations ^1–5^. Despite this growing recognition, our understanding of the landscape of somatic mosaicism remains incomplete, and its contribution to cellular phenotypic diversity and disease susceptibility remains unclear. There are two main obstacles to a better understanding of the functional implications of genomic mosaicism: amplification related biases and error in current genomic analysis methods for the discovery of mutations in single cells and failure to simultaneously capture aspects of the cell’s phenotype. Traditional bulk sequencing approaches, even when employing ultra-precise methods such as duplex sequencing or NanoSeq, average variant signals across heterogeneous cell types. The averaging masks cell–type–specific mutation burden and obscures sharing of mutations across cell lineages. In contrast, single-cell whole-genome amplification (scWGA) techniques such as the primary template–directed amplification (PTA) approach enable the discovery of the constellation of mutations in a cell independently of their overall frequency in tissue, with relatively uniform genome coverage, approaching that of bulk sequencing ^6–9^.

Until recently, scWGA did not allow the isolation of RNA from the same single cell, impeding both cell type identification and a direct readout of the potential consequences of sequence variants found on the DNA level. To bridge this critical gap, we benchmarked the ResolveOme method in human cells for parallel analysis of genomes and transcriptomes at single-cell resolution. ResolveOme extends the high-fidelity benefits of PTA by incorporating an initial reverse transcription step capturing polyadenylated mRNAs before genomic DNA amplification from a single cell (BioSkyrb genomics, USA). By comparing somatic mutations discovered from ResolveOme with standalone PTA and induced pluripotent stem cell (iPSC) lines derived from single cells of the same tissue, we have established its sensitivity, accuracy, and applicability to diverse sample types. scWGA can provide a temporal and mechanistic view of somatic evolution by using endogenous somatic mutations as intrinsic barcodes to map mitotic ancestry, time the origin of variants, and quantify clonal expansion dynamics.

Finally, to explore the capability of scWGA technologies to discover structural variations (SVs), we used an orthogonal amplification-free technique, Strand-seq, for high-throughput single-cell analysis of 200 kbp and larger SVs, including copy-number neutral events, in the same primary sample vis-a-vis scWGA data. This study provides a multidimensional framework integrating clonal iPSC lines with extensive scWGA and Strand-Seq datasets, and quantifying sensitivity, specificity, allelic dropout, and coverage uniformity which ensures accurate somatic mutation profiling. Our comparative analysis of the 3 tissues across all techniques enabled an analysis of differential susceptibility to mutagenesis across cell types and an accurate reconstruction of cell lineage phylogenies. The significance of this work lies in its potential to transform our ability to interpret somatic mosaicism within a functional framework.

## Results

### Study overview

To benchmark and validate the scWGA protocol against an orthogonal clonal approach, we leveraged a panel of 35 clonal iPSC lines, derived from eight body locations of a healthy donor, NC0 (**Fig. 1A** and **B**). The iPSC lines, by virtue of their clonal expansion, provide a high-confidence somatic variant truth set free of amplification artifacts, enabling us to assess whether single nucleotide variants (SNVs) identified in single-cell PTA or ResolveOme libraries recapitulate bona fide mutations present in matched iPSC lines derived from the same biopsy. Among those, 13 iPSC lines were derived from NC0’s peripheral blood mononuclear cells (PBMC) using Sendai virus reprogramming, 8 from urinary tract derived cells, using an episomal transfection method, and 14 were previously generated from fibroblasts isolated from six primary skin biopsy sites (left arm, right arm, right thigh and left thigh, respectively LA1/2, RA1/2, RT and LT) (**Fig. 1A**) ^10^. All iPSC lines were expanded and sequenced by a standard short-read Illumina platform (mean depth ∼35X), providing a reference for somatic variant discovery (**Table S1**). To distinguish somatic mutations from inherited variants, we employed a tumor–normal variant-calling strategy using matched bulk blood DNA sequence data as normal reference (**Methods**). To ascertain that iPSC lines were clonal, we analyzed somatic SNVs in each line. In all cases, allele frequency distributions followed a unimodal, approximately Gaussian shape centered at around 0.5, confirming that all iPSC lines were monoclonal (**Fig. S1**).

**Figure 1:**
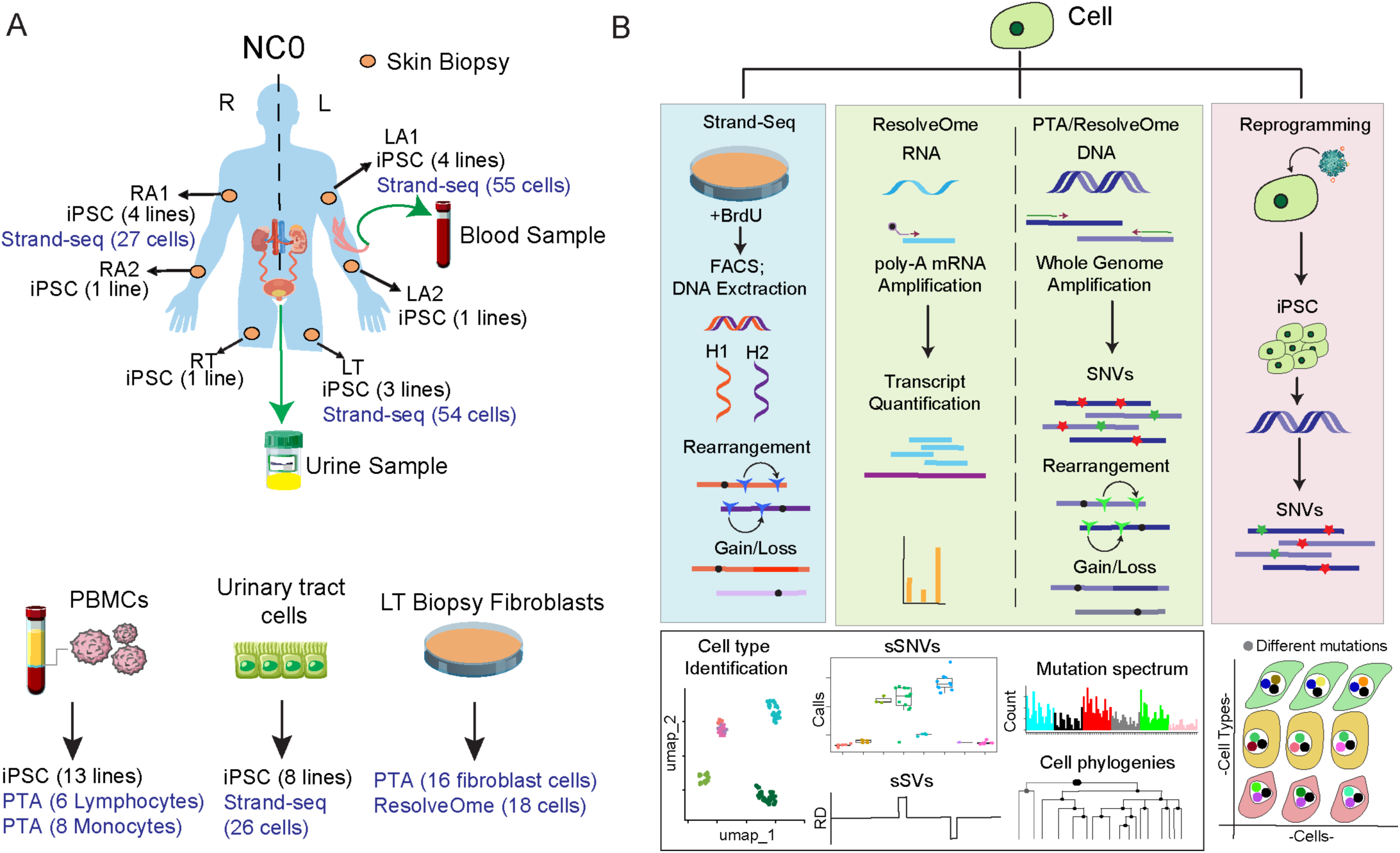
Study design and multi-platform workflow for profiling somatic mosaicism. **A)** Overview of sample collection for donor NC0. Biospecimens included skin biopsies (RA1, RA2, LA1, LA2, RT, LT), peripheral blood mononuclear cells (PBMCs), and urine-derived epithelial cells. These samples were used to derive 35 clonal iPSC lines and primary fibroblast cultures. **B)** Integrated analytical workflow depicting scWGA by the PTA or ResolveOme approach, generation of iPSC lines, and Strand-seq for haplotype-resolved structural variant detection. Downstream analyses included transcript quantification, SNV calling, CNV and SV analyses, mutational signature deconvolution, and reconstruction of cell-lineage phylogenies.

We next applied scWGA approaches, both standalone PTA and as part of ResolveOme, to fibroblasts derived from the LT biopsy (n=34 cells) and to lymphocytes and monocytes isolated and identified by FACS from PBMCs (n=14 cells) of the NC0 subject (**Fig. 1A** and **B**). For LT fibroblasts, cells and nuclei were sorted by FACS to isolate single cell/nuclei and carry out scWGA (**Methods**). As the standard ResolveOme protocol resulted in low RNA yield in nuclei samples (**Table S2; Methods**), we optimized the standard ResolveOme workflow by modifying sort-buffer volume and bead-cleanup steps, and then used this modified protocol to co-amplify genome and transcriptome. Lastly, we applied Strand-Seq to fibroblasts and urinary tract cells, leveraging its haplotype-resolved readout on the single cell level (**Fig. 1B**).

### Cell type identification by single-cell level transcriptome profiling

We evaluated the ability of ResolveOme to identify cell type from single-cell transcriptomes by comparing whole cell and nuclear preparations derived from the 18 fibroblasts derived from the LT biopsy. Transcriptome libraries were prepared as follows: (i) for cells with the standard protocol (n=6), (ii) for cells with a modified protocol (**Methods**) (n=8), (iii) for nuclei with the modified protocol (n=10). In addition, a bulk RNA-seq library was prepared from the parental LT fibroblast culture from which the WGA-cells and nuclei were derived. The obtained transcriptome data was processed with identical preprocessing, normalization, and dimensionality-reduction pipelines and then integrated with a reference dataset we had produced earlier ^11^, which consists of bulk RNA-seq profiles of fibroblasts, iPSCs, and induced neurons. The comparison with the reference dataset revealed that fibroblast cells processed with standard or modified protocols, as well as nuclei processed with the modified ResolveOme protocol, clustered tightly with the reference fibroblast populations, indicating that the approach can preserve canonical transcriptional signatures and enables accurate cell-type classification (**Fig. 2A**). The bulk RNA-seq sample from the primary LT fibroblasts also clustered coherently, further confirming the consistency and robustness of fibroblast identity. Reference cell-type populations formed non-overlapping clusters, serving as effective controls for cell-type identity.

**Figure 2.**
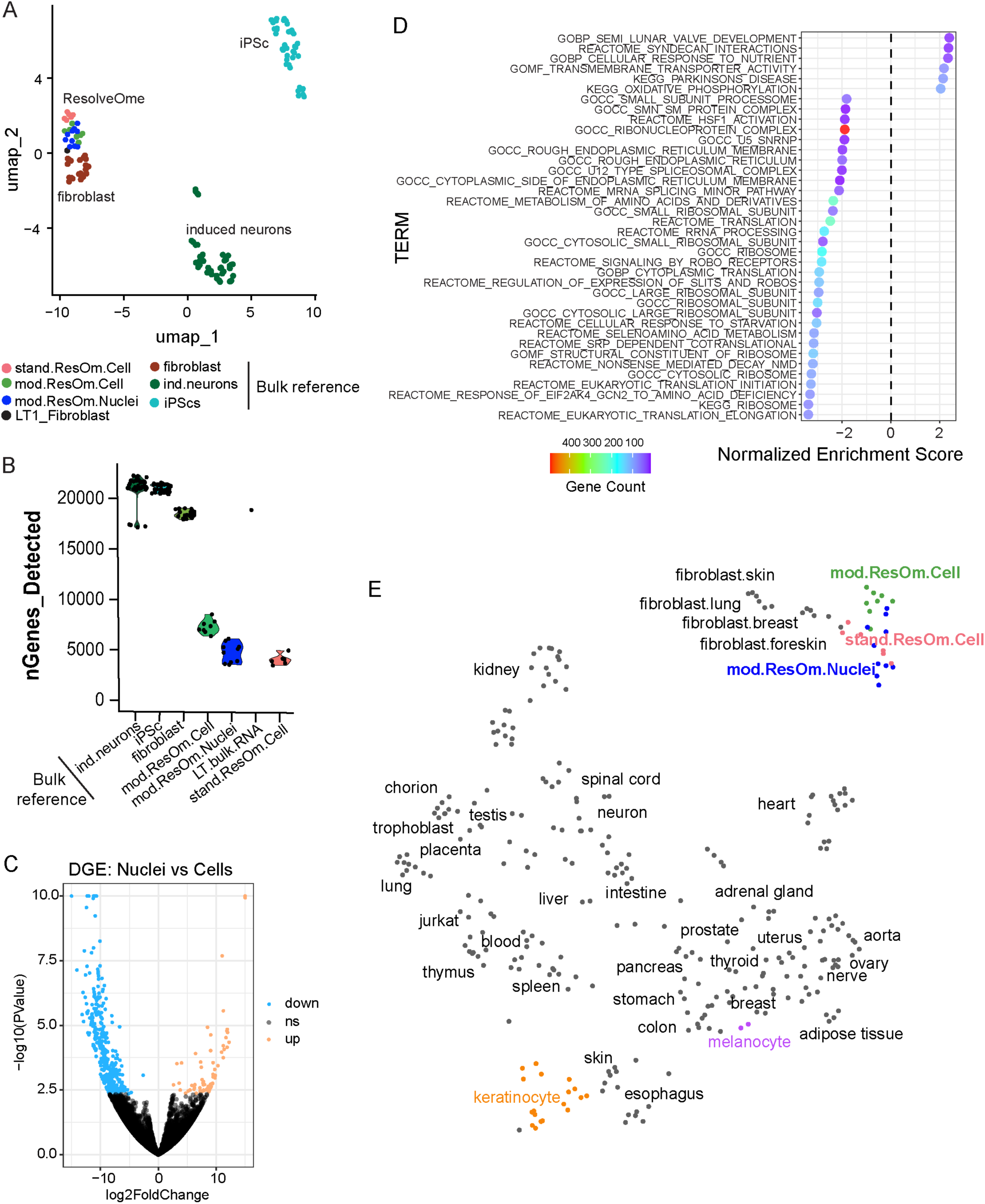
Identification cell type from ResolveOme single cell RNA-seq profiles. **A**) UMAP plot of RNA-seq data from different samples. Each dot represents a bulk sample, a cell or a nucleus. **B**) Number of detected genes in bulk RNA-seq data from reference fibroblast, iPSC, and induced neurons samples as compared to single cell transcriptomes in cells and nuclei. Each dot represents an individual cell or bulk sample. **C)** Volcano plot of differential gene expression in fibroblast nuclei vs cells. Genes with significant downregulation (blue; log2FC < −1, adj. p < 0.05) and significant upregulation (orange; log2FC > 1, adj. p < 0.05) are highlighted, while non-significant genes are shown in grey. **D)** Gene set enrichment analysis for transcriptome data from nuclei and cells (FDR < 0.05). Datapoints with negative x-axis values highlights decreased enrichment of gene ontology or pathway in nuclei transcriptomes and vice versa. Colour gradient reflects the count of genes associated with each term (blue = low count; red = high count). **E**) Projection of the transcriptome data from ResolveOme-amplified single fibroblast cells or nuclei (red, cell standard protocol; green, cell and blue, nuclei, modified protocol) to a UMAP representation of transcriptomic data from a broad range of cell types from Boix et al. (2021).

Since we did not obtain enough RNA yield from nuclei processed with the standard ResolveOme protocol, these samples were not processed for libraries or sequenced. However, the modifications in the ResolveOme protocol allowed comparable gene detection (**Fig. 2B**) and RNA yields (**Table S2**) in cells and nuclei. Comparison of gene expression profiles between cells and nuclei revealed that ∼450 genes were downregulated and ∼150 genes were upregulated in the nuclei samples (**Fig. 2C; Table S2**). Functional enrichment analysis of the downregulated gene set highlighted systematic depletion of transcripts related to cytosolic processes, including translation, ribosome assembly, and protein synthesis (**Fig. 2D; Table S2**). Most of the underrepresented genes in nuclei samples encoded ribosomal proteins and other components of the translational and splicing machinery, consistent with the expected loss of cytoplasmic content in nuclear preparations.

To further evaluate the performance of the ResolveOme protocol in distinguishing cell types across broader tissue contexts, we projected our data onto the single-cell reference atlas from Boix et al. ^12^, which includes diverse primary human cell populations and resolves the identity of cell types despite their close developmental origin (**Fig. 2E**). ResolveOme-amplified samples, both cell and nuclei, aligned with reference fibroblast clusters derived from skin, breast, and lung tissues and remained transcriptionally distinct from related cell types from dermal origin such as keratinocytes and melanocytes (orange and purple, **Fig. 2E**). Despite their shared dermal origin, each of these cell types formed clearly separated clusters, reinforcing the specificity of cell-type resolution in the integrated UMAP. Together, these results demonstrate that the modified ResolveOme protocol allows for recovery and amplification of single cell RNA from nuclear preparations, which maintains high-resolution transcriptional fidelity comparable to RNA isolated from cells, enabling robust and accurate cell-type identification.

### Single-cell somatic SNV discovery reveals somatic mutation heterogeneity in tissues

To assess the somatic variants across different biopsies, we analysed scWGA and iPSC’s whole genome sequencing data. We performed a rigorous, two-stage quality control (QC) workflow to assess whole genome amplification by PTA and ResolveOme on single cells and nuclei. In the first QC tier, all 48 samples (34 from LT fibroblasts and 14 from PBMC) underwent a multiplex 4-loci PCR assay to verify successful whole-genome amplification. Samples that generated all four expected amplicons were advanced to sequencing. In the second QC tier, we sequenced the scWGA samples at 35x genome coverage and subjected WGS data to the SCELLECTOR pipeline for evaluation of allelic balance at germline single-nucleotide variants. Only scWGA samples whose variant allele frequency (VAF) histograms exhibited a tight, bell-shaped distribution centered around 50% (standard deviation ≤ 0.23) and allele drop rate <8% were deemed to pass (**Fig. 3A & S2**). By this criterion, 39 WGA samples out of 48 (81.25%) met our stringent QC thresholds, demonstrating minimal allele dropout and amplification bias. By QC measures, all ResolveOme samples were virtually indistinguishable from standalone PTA. These combined QC measures confirm that our optimized ResolveOme protocol reliably generates high-quality DNA libraries from both cells and nuclei.

**Figure 3.**
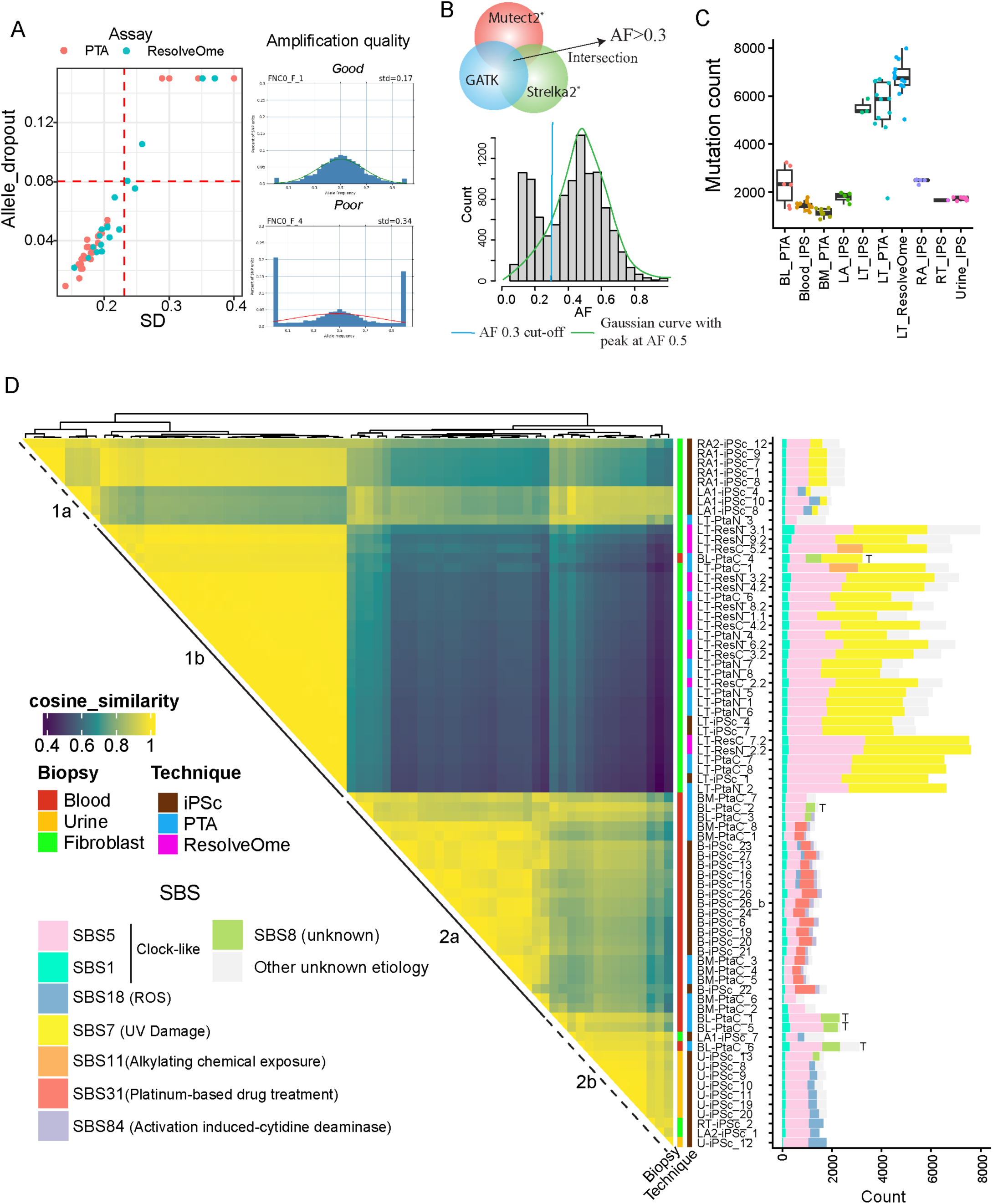
Mutational spectra and signature contributions across single cells and clonal lines. **A)** Scatter plot of SCELLECTOR allele-dropout assessment for PTA (red) and ResolveOme (blue) libraries. Samples below the red dashed thresholds were deemed QC-pass for downstream analysis. Examples of germline SNPs scored via SCELLECTOR are shown on the right. **B)** Strategy for calling somatic mutations. Inset: bar plot of sSNVs VAF in a cell at 0.3 cut-off. **C)** sSNVs mutation burden across different samples from different biopsies. **D)** Mutation spectrum similarities and SBS signature profiling of sSNVs called from WGA method and clonal iPSC libraries. Heatmap reflects cosine similarity computed from 96 trinucleotide motif. Sample labels are color-coded by iPSC origin or amplification approach. Right panel: stacked bar plots show SBS signature decomposition and sSNV burden per sample. Sample code: LA, RA, LT, RT, B/BL/BM, U, indicate sample origin as shown in Fig.1 (left arm, right arm, left thigh, and right thigh biopsies, blood, and urine); iPSC, PtaC/PtaN, ResC/ResN indicate iPSC clone, PTA cell or nucleus, ResolveOme cell or nucleus, respectively); and last digits sample number. T on bar plot represents T-lymphocytes (see Fig. 5D).

We next called somatic SNV (sSNV) across both scWGA and iPSC genome data by applying a three-caller consensus approach for Mutect2, GATK HaplotypeCaller, and Strelka2 (Mutect2 ∩ GATK ∩ Strelka2) and imposing an allele-frequency (AF) minimum threshold of 30% to exclude subclonal or artefactual calls since bona fide somatic heterozygous SNV calls have an AF distribution close to 50% in single cells or clones (**Fig. 3B; Methods**). This approach yielded a remarkably consistent mutation burden in each biopsy site with little variation between technology used – PTA, ResolveOme or clonal iPSC lines. An overall higher sSNV burden was detected in fibroblasts from the LT biopsy, with an average of ∼6,500 sSNVs per cell in scWGA samples and a comparable burden of 5,575 ± 325 sSNVs in 3 iPSC lines from the same biopsy. This finding is consistent with mutation heterogeneity across cells within individuals (**Fig. 3C**).

We next clustered scWGA samples and iPSC lines by similarity in mutation spectra (i.e., distribution of sSNVs in trinucleotide motifs) to reveal that, with a few notable exceptions discussed below, single-cells co-clustered with iPSCs derived from the same biopsy/cell type, demonstrating that technical variance from amplification and library preparation is far outweighed by genuine biological differences between cell types and cells (**Fig. 3D**). To put our analysis in the context of known mutation processes, we also decomposed mutation spectrum in each sample and line into known mutational signatures from the COSMIC database. Across all samples, clock-like signatures SBS1 and SBS5 together account for ∼40–60% of sSNVs, reflecting universal endogenous deamination and methylation damage (**Fig. S3**). Rare and low-level contributions to mutation spectrum (typically <20%) from assay-specific or stress-related signatures such as oxidative damage (SBS18), damage by alkylating agents (SBS11) and other signatures of unknown etiology likely represented residual amplification artifacts or inaccuracies in signature decomposition.

Overall, iPSCs/scWGA-samples clustered into two major clades (**Fig. 3D**). One large cluster consisted of fibroblast-derived iPSCs/scWGA-samples from the four skin biopsies and was characterized by a contribution of UV-induced mutations (SBS7). This major cluster was sub-divided in 2 sub-clusters distinguished by the fraction of UV-induced mutations. The smaller subcluster (1a) consisted of fibroblast-derived iPSCs and was charactered by a smaller amount of UV-damage, while the bigger cluster (1b) contained samples with higher UV-damage (>50%) and was almost entirely composed of cells from the LT biopsy (**Fig. S3**). In most skin fibroblast-derived iPSC and WGA-cells from all biopsies, UV-associated signature SBS7 contributed ∼50% of mutation burden, consistent with lifelong sun exposure of dermal fibroblasts. However, the existence of two UV-damage associated subclusters points to heterogeneity of UV-associated mutagenesis in cells from various physical body locations. The ultimate manifestation of this heterogeneity was three fibroblast-derived iPSCs that had little or no UV-damage and clustered with iPSCs/cells from urine and blood that had no mutation from sun exposure.

The second major cluster was characterized by mutation spectra driven principally by endogenous, clock-like processes, chemotherapy or ROS signature. This cluster mostly consisted of cells and iPSCs from blood and urine segregated in subclusters 2a and 2b respectively. Urine derived iPSC lines were slow growing which corelates mutations accumulated due to prolong cell culture resulting in SBS18 signature. Contrary to fibroblasts, almost all blood- and urine-derived iPSCs/WGA-cells had no UV damage. But one blood-derived PTA WGA-cell (BL-PtaC_4) had high mutation burden and high fraction of UV-induced mutations, so that it clustered with UV-affected iPSCs and fibroblast cells (**Fig. 3D**). This cell was deemed to be a T-cell based on rearrangements in T-cell receptor (TCR) loci (see next section), so its UV exposure can be attributed to residing in or traveling through dermis for a significant amount of time, and thereby being exposed to UV-damage ^13^. Hence, mutational signature likely reveals the history of a cell.

Blood derived iPSCs/WGA-cells formed a subcluster (2a), which is likely attributable to the contribution of signature SBS31 to their mutation spectrum. That signature is hypothesized in COSMIC to originate from chemotherapy, consistent with the donor having a history of chemotherapy and the signature being observed only in the blood cells. Interestingly, all lymphocytes and some monocytes didn’t have detectable contribution from SBS31, possibly implying heterogeneous effect of chemotherapy in blood, similar to the heterogeneous UV-damage signature in the skin. Finally, among blood-derived cells, five WGA-cells, identified as T-cell based on rearrangements in TCR loci (see next section), had about twice the number of mutations compared to monocytes and iPSC lines (**Fig 3C**). The increase in burden was because of higher count of mutations from signature SBS8 (unknown etiology) and for one cell (as mentioned above), from UV damage. This observation is consistent with T-cells having long lifespan and matches the mutation profile for B- and T-cells ^14^. Intermixing of single cells and iPSC lines in clusters and the concordance of their mutation spectra establishes PTA and ResolveOme as robust platforms for accurately capturing somatic mutation landscapes across diverse tissues and conditions.

### Single cells enable accurate reconstruction of cell phylogeny

Cell phylogenies in NC0 were previously studied by sequencing 14 fibroblast-derived iPSC lines and utilizing shared mutations as markers for the most recent common ancestral cells^10^. That analysis revealed sharply asymmetric contributions of the first two post-zygotic lineages to tissues (i.e., in blood 90:10 and urine 70:30), leading us to annotating those lineages as dominant and recessive. Here, we extended the previous data with new sequencing data from PTA- and ResolveOme-amplified PBMCs and fibroblasts and added new iPSC lines derived from PBMCs and urinary tract-derived cells, totalling 51 samples: 29 iPSC lines, 22 PTA/ResolveOme cells (**Table S1**).

There are three confounding factors when using single cell data for lineage reconstruction: (1) low-level amplification artefacts result in sharing of false positive mutations between cells; (2) dropouts in 5-10% of cells’ genomes result in germline variants falsely appearing as high-VAF mutations, i.e., mutations shared by most but not all cells; (3) somatic deletions and LOH events in cells may also lead to germline variants falsely appearing as high-VAF mutations (**Fig. 4A**). The latter two factors are of particular concern because, as mentioned above, mutations from the dominant lineage in NC0 are present in 70-90% of cells and are hard to distinguish from germline variants, and such lineage asymmetries are common in the human population^15,16^. To generate the candidate set of shared somatic mutations we used the previously developed All^2^ pipeline which performs comprehensive pairwise comparisons across genomes of single entities (cells or clones) and computes germline and somatic scores for each mutation^17^. A mutation identified in a small fraction of cells has high somatic and low germline score, while a mutation identified in almost all cells has high germline and low somatic score. We then developed a set of filtering steps to eliminate falsely appearing shared and retaining true shared somatic mutations for tracing cell lineages (**Methods**). Briefly, in all entities, we identified large deletions and LOHs, and in PTA/ResolveOme cells we additionally identified regions of allele dropouts (**Fig. 4A**). For mutation within these regions, we used the allele drop-out mode in All^2^ to reduce counts of likely germline variants (**Fig. 4B, C**). For the same reason, we further removed calls that matched genomic variants frequently observed in the population or found in all but one entity. Next, we eliminated likely low-level amplification artifacts, i.e. calls with combined VAF across shared cells significantly below 50% (**Fig. 4D**). Next, using shared somatic calls (red line in **Fig. 4C**), we generated an initial scaffolding of lineage tree, queried the likely germline calls for consistency with branches in the tree scaffold and filtered out calls that were inconsistent (**Fig. 4E**). All these filtering steps reduced the number of shared mutations from 128,340 to 51,109 (**Fig. 4F, S4B**). Remarkably, the number of calls in the “likely germline” category were reduced from 18,576 to just two calls, of which one was identified previously as a true somatic mutation (denoted as ‘a’, **Fig. S4A**), while the other one is likely a germline SNP.

**Figure 4.**
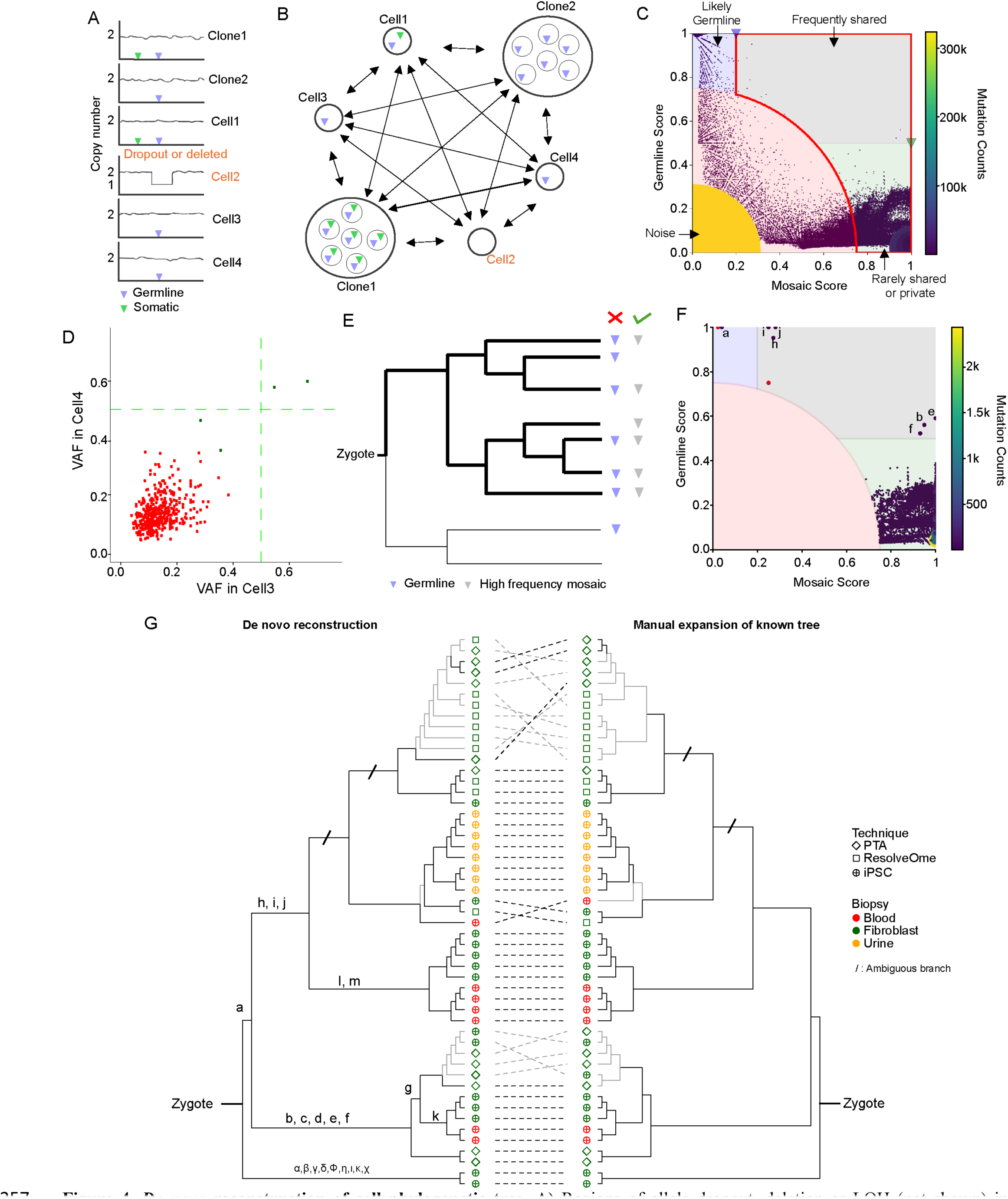
*De novo* reconstruction of cell phylogenetic tree. **A)** Regions of allele dropout, deletion or LOH (not shown) in individual cells will interfere with lineage reconstruction and need to be identified and accounted for. **B)** Comparing all-to-all entities (cells or clones) calls mutation candidate with associated germline and somatic scores. **C)** The two scores categorize mutation calls (dots) into four groups. The red lines outlines calls used for the initial tree scaffolding. **D)** Many shared calls between two cells have VAF significantly below 50% in both cells and were filtered out. **E)** Initial tree scaffolding based on shared mutation calls and the concept of filtering likely germline variants. **F)** Final mutation call sets across four categories. Previously known embryonic mutations are annotated with letters. Two likely false positive mutations are shown in red. **G)** Comparison of lineage reconstruction by the *de novo* approach developed here and manual expansion of the previously known tree^10^. Branches with the same (black) and different (grey) topologies are annotated.

Using all calls that passed filtering, we reconstructed the final lineage distribution using the maximum parsimony principle implemented by MPBoot (**Fig. 4G, S5A**). When comparing this *de novo* lineage reconstruction with manual expansion of the previously reported tree^10^ (**Methods**), we found that all early branches were correctly reconstructed while there were a few discordances in the later branches involving PTA/ResolveOme cells. This is expected, as earlier branches share mutations across multiple cells, and, with germline contaminating variants removed, those shared mutations make tree branching unambiguous for *de novo* reconstruction and manual expansion. On the contrary, later branches share mutations across a few cells and in some cases, ambiguity can arise due to noise from amplification. Given that the manual tree expansion does not account for amplification noise during WGA and allele dropouts introduced by uneven amplification, it is likely that branching in the *de novo* tree is more accurate.

In comparison to the previously known tree, the *de novo* reconstructed tree revealed more resolved and later branches, consistent with analysing more cells than previously (**Fig. S5B, Methods**). Interestingly, almost all resolved late clades (separated from the zygote by at least 500 mutations) likely represented clonal expansions as evidenced by missense or truncating SNVs in known cancer driver genes from Cancer Gene Census Tier-1 (SETBP1, NF1, FLT3, TRIP11, PTPRT, ATR, MECOM, ATP2B3, NCOR2, ROS1, FGFR1, and MYH9) (**Fig. S5C**). In conclusion, we demonstrate that removing amplification noise and allele drop-out during scWGA allows for a combined analysis of shared mutations found in iPSCs and WGA-single-cells, enabling precise and complete cell lineage reconstruction.

### Large aneuploidies and smaller copy number alterations at single cell resolution

To analyse aneuploidies and copy number alterations (CNAs) in the fibroblast- and blood-derived PTA- and ResolveOme-amplified cells (n=39), we performed read-depth segmentation on 100kb bins using CNVpytor^18^. In fibroblast scWGA samples, 3 out of 25 cells/nuclei (two ResolveOme, one PTA) revealed a chr21 gain, and 2 PTA cells carried X monosomy (**Fig. 5A**, **S6A**). Apart from these frequent events, we were able to detect singleton (i.e., in just one cell) alterations: a chr20 trisomy and a chr15 monosomy in fibroblasts and monosomy of chrX in a monocyte (**Fig. 5A, S6A**). Strand-seq analyses confirmed the frequent events, providing a better estimate for the frequency (6% and 32% for chr X monosomy and chr21 trisomy, respectively) of each aneuploidy in the LT fibroblast population (**Fig. 5B, C**). Further, Strand-seq confirmed two aneuploidies detected in iPSCs from the LA biopsy (**Fig. 5B**).

**Figure 5.**
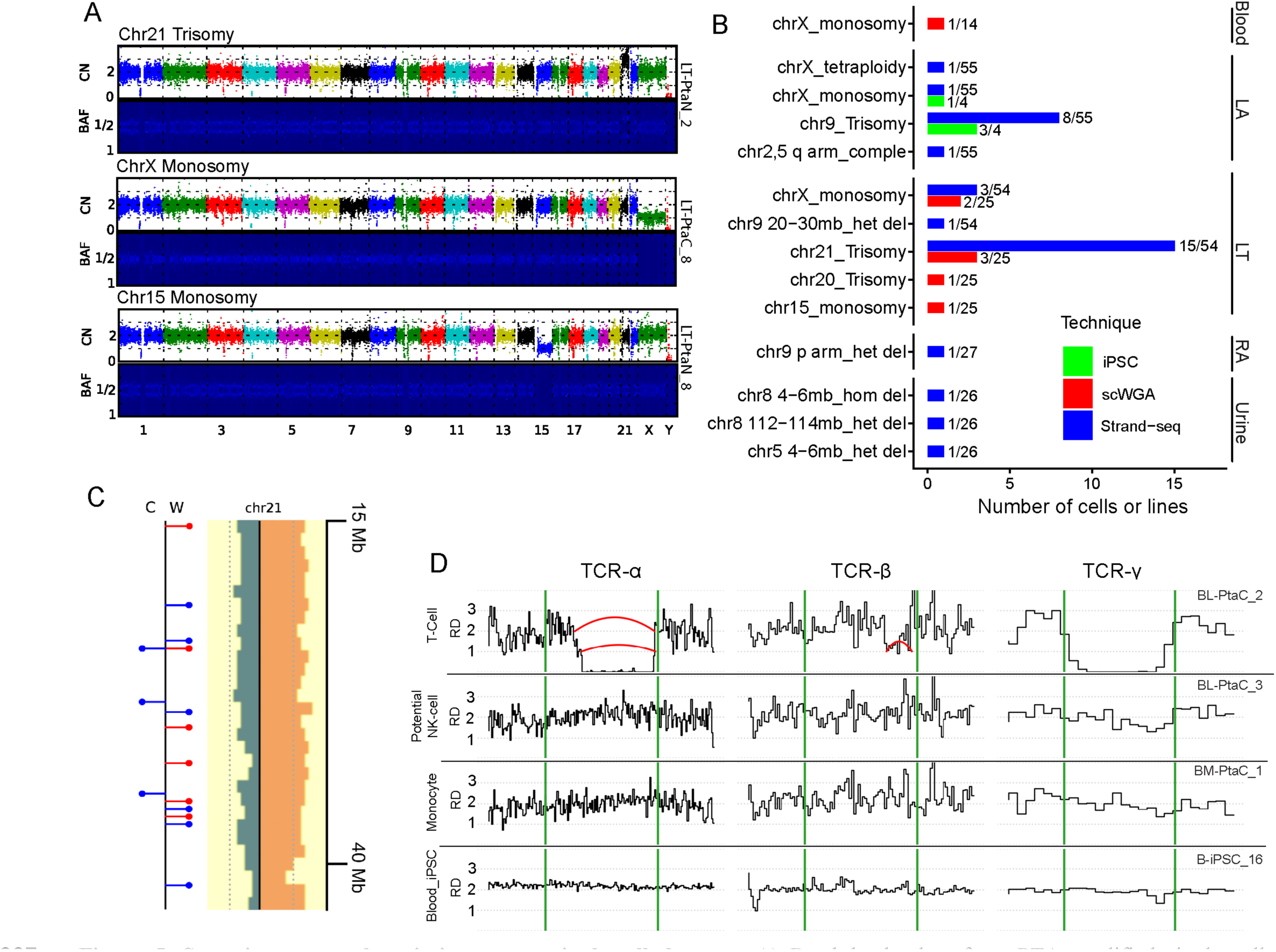
Somatic structural variation across single-cell datasets. **A)** Read-depth plots from PTA-amplified single cells highlighting copy-number variation: chr21 trisomy, chrX monosomy and chr15 monosomy. **B)** Summary of somatic structural variants (sSVs) detected across different biopsies from donor NC0. Fractions represent the number of cells or iPSC lines carrying sSVs versus the total number of cells/iPSC lines analyzed. **C)** Example of chr21 trisomy detected by Strand-seq in a fibroblast from LT biopsy. Read depth profile on Watson (W, orange) and Crick (C, teal) strands of chr21 long arm. A 2:1 ratio of W to C read depth is indicative of a trisomy. Positions of individual heterozygous SNPs of haplotype 1 (H1) and haplotype 2 (H2) are denoted as red and blue lollipops. SNPs on the Crick strand are shown on the left and those on the Watson strand on the right, with H2 SNPs present on both a Crick and a Watson copy, indicating duplication of the chromosomal homolog corresponding to H2. **D)** Copy number profile inferred from depth of coverage in TCR loci for 3 cells and iPSC from PBMC. Red arcs show deletions detected by discordant or split reads.

Additionally, we catalogued six classes of large somatic structural variants discovered by Strand-seq in 162 cells, including 54 LT fibroblasts — trisomy, tetraploidy, monosomy, homozygous deletion, heterozygous deletion, and complex rearrangements—across eight chromosomes. Most of these variants were present in one cell per biopsy site (likely representing cell-private events). Haplotype-resolved SNP phasing from Strand-seq provided additional validation and insight into the allelic origin of these events (**Fig. 5C; Methods**). We inferred duplication of haplotype 2 (H2) in LT cells with chr21 trisomy and loss of H2 in cells with X monosomy. To confirm the identity of SV-affected haplotypes, we compared haplotype-resolved SNP calls from Strand-seq with allele frequency of variants in the PTA data. Heterozygous SNPs of H2 on chr21 were consistent with variants of AF above 0.5 in the PTA cells and monosomic PTA cells retained hetSNPs that matched exclusively to haplotype 1 (H1) calls in Strand-seq data (**Fig. S6B, C**).

To test if smaller CNAs could be detected in scWGA-cells, we examined WGS data for PTA-amplified lymphocytes (n=6), monocytes (n=8) and PBMC-derived iPSC lines. In this targeted assay the majority of lymphocytes (5 of 6 cells) showed clear evidence of locus-restricted deletions at T-cell receptor loci (TCR). Typically, each deletion was accompanied by matching split reads or read pairs and by step-wise changes in local copy-number profile, all consistent with somatic V(D)J immune-rearrangement events (**Fig. 5D, S7**). By contrast, none of the monocytes and iPSC lines analysed exhibited rearrangements or deletions in TCRs, supporting their expected myeloid identity (**Fig. S8**). Taken together these observations validated scWGA to detect focal, biologically expected rearrangements at single-cell resolution and provided orthogonal evidence that iPSCs in this study derive from non-rearranged myeloid cells. Interestingly, one lymphocyte lacked detectable structural changes in TCRs suggesting that it is a natural killer (NK) cell (a type of lymphoid cell lacking V(D)J recombination), consistent with ∼10% of nucleated blood cells being NKs (**Fig. 5D, S7**).

## Discussion

This study provides a multidimensional framework integrating clonal iPSC lines with extensive scWGA and Strand-seq datasets, quantifying sensitivity, specificity, allelic dropout, and coverage uniformity which ensure accurate somatic mutation profiling. Single cell technologies based on *in vitro* whole genome amplification presented seemingly insurmountable drawbacks for somatic mutation discovery, mostly due to polymerase incorporation errors, uneven genome amplification and allelic drop-outs^19–21^. The current results demonstrate that genome amplification is remarkably accurate with PTA and ResolveOme technologies, and that the remaining errors can be efficiently filtered out bioinformatically, as shown by the concordance between PTA/ResolveOme and non-WGA technologies such as iPSC and Strand-seq in SNVs and SVs discovery. The resulting single cell-based datasets including single fibroblasts, blood-derived lymphocytes and monocytes and iPSC lines permitted a direct comparison of mutation burden and spectra across cell types, capturing different susceptibility to mutagenesis among cells, and allowing to infer aspects of cellular history (environmental exposure, migration, tissue residence) from patterns of cellular mutagenesis. scWGA also enabled the analysis of chromosomal aneuploidies and smaller SVs as in the case of V(d)J recombination in T-cells, suggesting that PTA and ResolveOme are comprehensive approaches for genome characterization in a cell.

Among the single cell whole genome discovery approaches, ResolveOme enables the combined discovery of single cell genomes and transcriptomes, which at the minimum reveals the identity of the cells carrying somatic mutations and potentially may reveal their phenotypic effects. Hence, these approaches can transform our ability to understand the significance of somatic mosaicism within a functional framework and to elucidate the logic of competitive phenomena among cells—for instance, linking a specific mutation with changes in gene expression and clonal expansion of a cell type. Additionally, both PTA and ResolveOme currently represent the most widely applicable way to map the specific constellation of post-zygotic mutations that is carried by each cell. This information enables the retrospective reconstruction of cellular phylogenies by assessing shared mutations across cells, which by definition are generated by their most recent common ancestor^10^. Mapping cell lineages uncovers cellular dynamics across development and aging, including positive and negative selection among cells leading to re-shaping of tissue’s composition during an individual’s lifetime, along with differences in somatic mutation distribution/inheritance across cells and tissues. Transcriptome characterization of the terminal cells in the lineage trees permit accurate coupling of lineage divergence with cell phenotypes.

In summary, these technologies will deliver a powerful resource for the scientific community enabling the precise cell-type assignment of somatic variants and the ability to draw lineage hierarchies, uncovering functional consequences of mosaic mutations, and paving the way for mechanistic insights into development, aging, and disease processes.

## Limitations of the Study

As shown here, the ResolveOme technology is a major step towards defining type and studying somatic mutations in a cell, yet the wide applicability of this technology across a variety of cells, tissues, and cells/tissue storage conditions remains to be established. A parallel study demonstrates applicability of PTA to analysis of single cells in post-mortem tissues (SMaHT package #5, “A comprehensive view of somatic mosaicism by single-cell DNA analysis”, submitted). Artifacts of amplification including errors in DNA sequence, uneven coverage, and allele drop-outs are currently inevitable and may hinder studies of cells with low mutation burden, which were not analysed here. The current study uses a single donor, and therefore the conclusions about heterogeneous susceptibility to mutagenesis across cells may not universally generalize across ages, exposures, or genetic backgrounds.

## Acknowledgements

We are grateful to the participant for the generous donation of samples. We thank the Yale Stem Cell Center for the generation of iPSC lines and the Yale Center for Genome Analysis (YGCA) for library preparation and sequencing. We acknowledge the Yale Center for Clinical Investigation for clinical support in obtaining the biopsy specimens and blood samples. We acknowledge the following grants support: NIH Common Fund SMaHT program UG3 NS132128 (AA) and UG3 NS132146 (AU); National Institute of Mental Health U01 MH106876 (FMV), and R01 MH131623 (AA). We also acknowledge the support of the National Institute of General Medical Sciences of the National Institutes of Health 1S10OD030363-01A1 (YCGA).

## Resource Availability

### Lead Contact

Requests for further information and resources should be directed to and will be fulfilled by the lead contact, Flora M. Vaccarino

### Material availability

Biological samples (iPSC lines, fibroblasts) will be available upon request, but we may require a completed materials transfer agreement and coverage of shipping costs. This study did not generate new unique reagents or DNA constructs.

### Data and code availability

This study was conducted as part of the NIH Common Fund consortium initiative, Somatic Mosaicism across Human Tissues (SMaHT). More information about the SMaHT Network and data sharing is available online at https://smaht.org and at https://data.smaht.org.

- De-identified human genomic sequence data will be deposited at dbGaP (accession number to be communicated) and will be made available under controlled access as of the date of publication. For reasons of subjects’ privacy, as specified in our consent form, we only allow for controlled access to genomic sequence data. In addition, processed datasets derived from these data will be available at dbGAP as of the date of publication.
- All original code is provided in this paper’s supplemental information.
- Any additional information required to reanalyze the data reported in this paper will be available from the lead contact upon request.

## Author’s Contributions

J.O.K., A.E.U., A.A., and F.M.V. conceived, directed, and supervised the study. A.N, R.P., L.T. and Y.H. performed cell culture. A.N., R.P., and Y.H. performed nuclear isolation and scWGA. R.P. developed the modified ResolveOme protocol and carried out ResolveOme data generation. E.B.G., and P.H. generated Strand-seq libraries. Y.J. performed sequencing data alignment for iPSC and scWGA samples. A.N., M.K.D., Y.J. and M.S. performed data driven quality check. A.N. conducted mutation analysis of PTA, ResolveOme, and iPSC WGS data, M.K.D. performed lineage reconstruction. M.S., D.O, and A.N conducted copy number and SV analyses. A.N., M.K.D., R.P., M.S., D.O., L.T., J.O.K, A.E.U, A.A., and F.M.V. wrote the manuscript.

## Methods

### Ethical considerations

Informed consent was obtained from the subject according to the regulations of the Institutional Review Board and the Yale Center for Clinical Investigation at Yale University.

### Fibroblasts isolation and fibroblast-derived iPSC generation

Small skin biopsies (3mm^3^) were collected from the inner area of the upper left and right arms (LA1 and RA1, respectively) and the front upper left and right thighs (LT and RT, respectively). Primary cultures of fibroblasts were derived from the biopsies using the standard explant procedure ^22^ and cultured as previously described ^23^. The fibroblast cell lines were reprogrammed into iPSC lines by a viral-free episomal reprogramming method using the Epi5 kit (Life Technologies) following the manufacturer recommendations. The iPSC lines were propagated using mTESR plus medium (StemCell Technologies) on dishes coated with Geltrex LDEV-Free Reduced Growth Factor Basement Membrane Matrix (Life Technologies) and propagated using Gentle Cell Dissociation Reagent (StemCell Technologies).

### Urinary tract cells isolation and urine cell-derived iPSC lines generation

Urine samples were collected in sterile urine collection cups using the midstream clean catch method. Cells from the urine samples were isolated and expanded following published protocols ^24,25^. Urine primary cell lines were reprogrammed into iPSC lines by a viral-free episomal reprogramming method using ReproTeSR (Stem Cell Technologies) and the Epi5 kit (Life Technologies) following the manufacturers recommendations. The iPSC lines were propagated following the same procedures as the fibroblast iPSC lines.

### Isolation of peripheral blood mononuclear cells (PBMCs) and generation of blood-derived iPSC lines

PBMCs were isolated from freshly collected blood using Histopaque 1077 (Sigma-Aldrich) following a standard density gradient procedure. Briefly, blood was collected using BD Vacutainer ACD tubes and immediately processed. After transferring to a centrifuge tube, the blood was diluted 1 in 3 with PBS, and Histopaque 1077 was gently pipetted at the bottom of the tube. After centrifugation, the mononuclear interphase was collected, rinsed with PBS and cryopreserved. iPSC lines were generated from the PBMCs by a Sendai reprogramming method using the Cytotune 2.0 kit (Life Technologies) following the manufacturer recommendations and were propagated using the same procedures as for fibroblast iPSC lines.

### Nuclei extraction

LT fresh fibroblast cells (∼2 million) suspended in media were pelleted (300xg 5mins, 4°C), washed twice with 5 ml PBS with 2% BSA to remove media, and resuspended in a final volume of 1.5 ml. The sample tube was split into 2 aliquots (2 x 700µl) containing approximately 1million cells per tube. Nuclei were extracted from one of the aliquots using the standard 10X Genomics protocols: (https://cdn.10xgenomics.com/image/upload/v1660261285/support-documents/CG000124_Demonstrated_Protocol_Nuclei_isolation_RevF.pdf), while the remaining aliquot with intact live cells was used for single cell FACS. Single cells and single nuclei were checked for viability with a Luna-FL fluorescent cell counter, cell counts (99% live), nuclei counts (96% dead) as expected.

### PTA FACS

The LT fibroblast line, peripheral blood mononuclear cells (PBMC) lymphocytes or monocytes were used for PTA experiments. For fibroblasts, cells/nuclei were stained with propidium iodide (PI) (100 ng/mL) for 5 min on ice, filtered through a 40 µm mesh, and loaded into a FACS sorter equipped with a 100 µm nozzle. For PBMCs, cells were sorted immediately after isolation using the Histopaque method. FSC/SSC were used to identify the main population for lymphocytes/ monocytes using standard criteria. Fibroblast live cells (PI^−^), or intact nuclei (PI^+^) were deposited as one event per well in single-cell mode.

### Standard ResolveOme FACS

LT fibroblast cells/nuclei were extracted as described above, filtered through a 40 µm flowmi tip strainer (SP Bel-art) using a wide bore tip, checked for viability (Luna-FL fluorescent cell counter) and counted before diluting to a final concentration of 200 cells/μl. Prior to loading on the FACS sorter, single cells were dual stained (active and passive) with Calcein AM (30nM final) and Propidium Iodide (100ng/mL). Single nuclei were stained with PI only. FACS collection was performed on a SONY SH-800 sorter instrument with a 130 µm nozzle gated to collect cell cycle phase singlets. For collection, a LoBind 96-well plate (Eppendorf) pre-aliquoted with 3μl cell buffer (BioSkryb genomics, ResolveOme kit) was set up on a pre-cooled platform. FSC/SSC were used to identify the main population, and live cells gated (PI^-^, Calcein AM^+^) or intact nuclei (PI^+^), then deposited as one event per well in single-cell mode.

### Modified ResolveOme FACS

The LT fibroblast line cells/nuclei were collected in 2μl Cell buffer (BioSkryb genomics, ResolveOme kit), all other steps were kept consistent with standard ResolveOme FACS. All PTA and ResolveOme FACS sorted sample plates were immediately tightly film-sealed and briefly spun down (500 xg) to collect the sample at the base of the well before freezing on dry ice. Plates were stored, frozen at –80°C until processed for scWGA protocols.

### Primary Template–Directed Amplification (PTA)

Single cells were sorted into 3µL of BioSkryb ResolveDNA Cell Buffer were used. PTA reactions were assembled by sequential addition of proprietary BioSkryb reagents. Amplified DNA was purified using the BioSkryb ResolveDNA Bead Purification Kit. DNA yield and fragment size (200–4,000 bp) were confirmed by Qubit HS dsDNA assay and Agilent TapeStation (HS D5000 ScreenTape).

### ResolveOme standard protocol

Standard BioSkyrb ResolveOme v1 protocols (BioSkryb Genomics, USA, TAS-036.3) were used, Briefly, sorted plates containing 3μl sample were taken from −80°C and thawed on ice. Reverse transcription was performed (42°C for 90mins, 50°C for 30mins) to prepare first-strand cDNA for all cytoplasmic RNA and any extra-nuclear (nuclear-membrane adherent or nuclear-pore localized) RNA present. For nuclei only extra-nuclear RNA transcripts would be available for this initial RT step. After completion of reverse transcription, nuclear lysis was performed at room temperature with OM2 reagent (20mins shaking at 1400rpm) on a thermomixer (Eppendorf). scWGA (30°C 10hrs, 65°C 3mins) was then performed on a Biorad thermocycler using the standard manufacturer protocols (Bioskryb genomics).

Affinity beads were then added to separate the two fractions (genome and transcriptome) with ‘first-strand cDNA’ binding to the magnetic beads and ‘amplified DNA’ fraction remaining in the supernatant. Amplified DNA (supernatant) was carefully transferred to a clean tube. On-bead cDNA amplification was performed post bead washes, by adding 25μl RNA amplification mix cycling 37°C for 15 mins, 95°C for 2mins followed by 24 cycles (95°C 15sec, 58°C 30sec, 68°C 4mins) and a final extension step at 72°C for 10mins. As noted in Bioskryb protocols, it is critical for the beads not to dry during the removal of the amplified DNA supernatant, therefore care was taken to ensure that 25μl RNA amplification mix was immediately added. Post amplification, all samples were bead purified and eluted in 42µl ResolveOme elution buffer. A quality check was performed here to determine recovery yields and fragment size (Qubit HS dsDNA kit and Agilent Bioanalyzer respectively) for both amplified cDNA and DNA fractions of each sample. Successful single cells/nuclei (with both fractions showing good yields) were moved forward to library preparation.

### Modified ResolveOme protocol

With the intent to make the nuclear membrane more porous or leaky, and capture some of the intra-nuclear transcripts for the initial RT reaction, single cell/nuclei were collected by FACS/FANS on a pre-cooled platform into 96-well Lo-bind plates containing 2µl cell buffer (Bioskryb genomics) in lieu of the 3µl volume in standard protocols. Plates were tightly film-sealed, briefly spun and frozen at −80°C as described above. Post-thaw of sorted sample plates on ice from −80°C, 1µl of fresh cold cell buffer was immediately added, followed by briefly centrifuging at 750xg in a pre-cooled centrifuge at 4°C to resuspend and gently agitate the freeze/thawed cell or nuclear membrane prior to the RT reaction. From initial thawing of sample plates until the completion of RT (post RT stop buffer addition), sample plates were consistently maintained on pre-cooled instruments rather than at room temperature as recommended (i.e. thermomixer for OME-RT reaction set at 8°C for mixing steps, and all centrifuge spins were performed at 4°C) to further preserve the RNA integrity for the modified protocol. One important factor at this point was to vortex well all reagents, including all enzyme tubes, in the kit before use. As bead steps were critical for recovery of genomic material one last modification, to minimize any loss of low frequency RNA transcripts from single nuclei, for all samples in the modified protocols group, the bead pellets were ‘air-dried’ on the magnet (during ethanol removal steps) instead of centrifuging beads for all bead wash steps. After these modifications to the initial collection and sample preparation all standard ResolveOme V1 manufacturer protocols (Biosykrb ResolveOme kit, TAS-036.3) were followed for RT, nuclear lysis, WGA with post amplification bead purified cDNA/DNA fractions eluted in 42µl ResolveOme elution buffer. A quality checkpoint was performed here to check recovery yields and size (qubit and bioanalyzer respectively) for both amplified cDNA and DNA fractions of the sample. Successful samples, with both fractions showing good yields and fragment sizes (based on manufacturer recommendations), were moved forward to library preparation.

### 4-Loci PCR Quality Control

Each scWGA product was screened by a multiplex 4-loci PCR assay to confirm uniform genome coverage. Primer sequences were as previously described^26^. Ten-microliter reactions contained 1 µL 10× PCR buffer, 0.6 µL 50 mM MgCl₂ (1.5 mM final), 0.5 µL 10 mM dNTP mix (400 µM each), 2 µL primer cocktail (0.5 µM each), 0.1 µL HotStarTaq DNA polymerase (0.5 U), template DNA (20 ng), and nuclease-free water to 10 µL. Thermocycling was performed on a Bio-Rad C1000 Touch: 94 °C for 15 min; 12 cycles of 94 °C for 1 min, 68 °C for 1 min (−1 °C per cycle), 72 °C for 1 min; 26 cycles of 92 °C for 1 min, 55 °C for 1 min, 72 °C for 1 min; and a final extension at 72 °C for 10 min. Successful amplification of all four loci was confirmed by electrophoresis on a 1.5% agarose gel; only QC-pass samples advanced to library preparation.

### Library Preparation and Sequencing

4 loci QC-pass PTA products (≥500 ng input) were converted into PCR-free Illumina libraries using the KAPA HyperPlus Library Prep Kit, following the manufacturer’s protocol. For all ResolveOme reactions, 100ng of purified DNA fraction and 20ng of purified cDNA was carried forward to the library preparation steps and standard protocols were followed as per BioSkryb library preparation kit protocols. PTA libraries were quantified by Qubit and TapeStation. pooled equimolarly, and sequenced on an Illumina NovaSeq X PLUS platform (150 bp paired-end reads, target 30× genome coverage per sample). For ResolveOme library sequencing, 4nM pool of all libraries was prepared combining each sample (genome and transcriptome libraries) at a 19:1 ratio DNA: RNA fractions, rather than a 4:1 ratio DNA:cDNA suggested by standard protocols. Sequencing (2×150bp) was performed on the Novogene 6000 instrument S4 chip, with a loading concentration of 1.8pM.

### Strand-seq Assay

Fibroblasts were cultured in DMEM+10%FBS+1%L-Glu+1%N.E. Aminoacids+1%P/S and urine epithelial cells were cultured in RE/MC proliferation medium + Factors. Strand-seq libraries were prepared using an automated liquid handling workstation (Biomek i7) according to an established protocol with volumes adapted to work in 96 well plates^27^. In brief, after recovery the cells were incubated with 40µM BrdU for one cell division (exact time varied depending on the sample, i.e. 24h) followed by crosslinking and bulk-MNase digestion of nuclei. Next, the population of cells that incorporated BrdU for a single cell division was sorted as single nuclei into 96 well plates using flow cytometry based on quenching of Hoechst by BrdU. De-crosslinking, protease digestion and ligation of Illumina adapters was done as described in the original protocol with adjustments to volumes used. Adapter dimers were removed using a bead-based cleanup at a 0.8x bead:DNA ratio. The removal of BrdU-substituted DNA strands was done by Hoechst and UV-light exposure. Finally, the libraries were PCR amplified for 15x cycles, individual cells were simultaneously barcoded using a dual indexing strategy with iTru adapters. After the bead-based cleanup at a 0.8x bead ratio and pooling, size-selected libraries were subjected to deep sequencing with the Illumina NextSeq500 platform (MID-mode, 75 bp paired-end protocol).

### Data processing

#### WGS alignment

Paired-end FASTQ files were aligned to GRCh38_noALT using BWA-MEM v0.7.17 with read-group tags (@RG \tID=<sample= \tSM=<sample= \tPL:ILLUMINA \tLB:lib1 \tPU:unit1). The SAM output was piped to Samtools v1.9 for conversion to BAM (samtools view -b), then sorted (samtools sort) and indexed (samtools index). To correct misalignments around small insertions and deletions, GATK v3.8 RealignerTargetCreator was run with known -indel resources (Mills and 1000G gold standard.indels.vcf and 1000G phase1.indels.vcf) to define target intervals. IndelRealigner (GATK v3.8) then performed local realignment of reads within those intervals, and the resulting BAM was reindexed. Base quality scores were recalibrated using GATK v4.1.9.0. First, BaseRecalibrator was executed on the realigned BAM with known-site VCFs (dbSNP build 138, Mills and 1000G gold standard.indels, 1000G phase1.indels) to produce a recalibration table. ApplyBQSR then applied the recalibration to generate a final BAM file, which was indexed for downstream variant calling. A high-confidence set of germline heterozygous SNPs was generated from bulk NC0 blood.

#### Strand-seq

Strand-seq raw sequencing data was processed with MosaiCatcher v2 ^28^. The pipeline performs read alignment to GRCh38 reference genome, binning, multi-step normalization of read counts per bin, calling of heterozygous SNVs with freebayes and their phasing with StrandPhaseR ^29^. Strand-Seq directional reads were then used to build chromosome-wide haplotype blocks (H1 and H2) via StrandPhaseR. SV calling was done using the in-house data postprocessing and visualisation python-based tool *strandtools*^30^. Briefly, normalized read counts were merged into 1 Mb bins, followed by prediction of copy number states for individual bins and segmentation of chromosomes based on copy number and changes in Watson(W)/Crick(C) counts ratio. Manual inspection of segments was done to identify and classify both germline and mosaic SVs. Aneuploidies were detected as chromosome-wide deviations from the expected diploid ratios of W/C reads, interstitial deletions appeared as segments of twofold or fourfold drop in coverage in case of heterozygous or homozygous deletions respectively. Complex chromosomal alterations had multiple consecutive segments of copy-number changes. For each SV, phased SNPs were extracted within the affected chromosome and compared their allele representation in PTA libraries.

#### Detection of focal copy-number variations in T-cell receptor loci

Somatic structural variants and focal copy-number variations were identified using Manta (v1.6.0) and CNVpytor (v1.3.2). Manta was executed with default parameters, using each single-cell genome as the test sample and the corresponding bulk blood DNA as the matched control to enable detection of somatic SVs. CNVpytor analyses were performed with a bin size of 10,000 bp, deriving read-depth (RD) signals directly from the alignment (BAM) files. Heterozygous single-nucleotide polymorphisms (SNPs) identified from bulk blood DNA were used to compute B-allele frequencies (BAFs) in single-cell data, based on read counts obtained with *samtools mpileup* from single-cell alignment files. Visualization of genomic alterations was performed by integrating RD and BAF profiles from CNVpytor with split-read and paired-end junctions identified by Manta. In five of six PTA-amplified lymphocytes, distinct locus-restricted deletions were detected within T-cell receptor loci, characterized by stepwise decreases in RD, loss of heterozygosity at informative SNPs, and frequently supporting junctions identified by Manta, consistent with canonical somatic V(D)J recombination events. In contrast, no rearrangements or copy-number changes were detected in the corresponding TCR loci of monocytes or iPSC lines.

#### Variant Calling and Filtering

##### Somatic Variant Calling with Mutect2 and Strelka

Mutect2 (GATK v4.1.9.0) was run in tumor–normal mode using the --tumor as single-cell sample BAM and bulk blood WGS BAM as normal. The reference was GRCh38_noALT, and common-site VCFs (gnomAD, dbSNP) were provided to Mutect2’s --germline-resource and --panel-of-normals options. The unfiltered somatic VCF (*.unfiltered.vcf.gz) was processed with gatk FilterMutectCalls using default contamination estimates and read orientation models to produce the filtered somatic ‘PASS’ call set. Similarly, Strelka2 v2.9.10 was configured for tumor–normal analysis via configureStrelkaSomaticWorkflow.py with the single cell samples as tumor, bulk blood BAMs and the GRCh38_noALT reference. Somatic SNV and indel ‘PASS’ calls were extracted from the resulting VCFs (somatic.snvs.vcf.gz, somatic.indels.vcf.gz) and kept without additional post-filtering beyond Strelka2’s built-in filters.

##### Variant Calling with GATK HaplotypeCaller and VQSR

GATK HaplotypeCaller was run in cohort mode to generate raw germline VCFs (*.raw.vcf.gz) for each sample against GRCh38_noALT. VariantRecalibrator was applied separately to SNVs and indels using truth/training sets (HapMap, Omni, Mills, 1000G) to build Gaussian mixture models. ApplyVQSR was then used at tranche sensitivities of 99.5% (SNVs) and 99.0% (indels) to produce high-confidence calls. Germline filtering using the BSMN germlinefilter.py script to remove any variant present in the VQSR-filtered calls at the same genomic position and allele, ensuring retention of true somatic events.

Somatic calls from Mutect2, Strelka2, and GATK were intersected by matching chromosome, position, reference, and alternate alleles. Only variants present in all call sets were advanced. From this intersected list, single-cell SNVs (sSNVs) were selected by applying an allele-fraction cutoff of ≥0.30. Fitting a Gaussian to the AF histogram allows us to place a conservative cutoff at AF = 0.30, thereby capturing true sSNVs present in at least 30% of reads and minimizing the impact of allelic dropout or amplification noise.

#### Cosine similarity and signature decomposition

For each sample trinucleotide motif counts were generated and signatures were decomposed based on COSMIC database v3.4 with SigProfilerAssignment^31^. Pairwise similarity between reconstructed 96-spectra profiles was quantified using lsa::cosine function. Similarity matrix was used for hierarchical clustering and heatmap visualization (using ComplexHeatmap) with annotation tracks for biopsy source and technology. Reconstructed profiles for each sample, after which closely related SBS subtypes were aggregated into biologically interpretable groups both absolute and percent stacked exposure plots. For plotting we combined SBS7 subtypes into SBS7, and binned signatures with unknown etiological origin into an “Unknown_etiology” category.

#### RNA-seq Alignment and Quantification

Raw paired-end FASTQ files were aligned to the human GRCh38 reference genome (GENCODE v34) using STAR v2.7.3a in two-pass mode with default settings, producing coordinate-sorted BAM outputs. Gene- and transcript-level abundances were then quantified directly from the STAR alignments with RSEM v1.3.3 (rsem-calculate-expression --paired-end --bam), generating per-sample expected counts. For each cohort (ResolveOme and external datasets), we merged individual RSEM count tables by gene identifier into a unified count matrix. These merged matrices served as the input for all downstream single-cell and bulk analyses.

Merged count matrices were loaded into R v4.1.2 as Seurat v4.1 objects. We performed initial quality control by excluding cells with fewer than 200 detected genes or cell lines. Data were log-normalized (scale.factor = 1e4), and the top 3,000 highly variable genes were identified using the “vst” method. We then scaled expression values followed by RunPCA. UMAP embeddings were computed for visualization.

#### Differential Expression Analysis

For comparisons between conditions (e.g., Nuclei vs. Cells), we imported RSEM expected counts into R using tximport (type = “rsem”, tx2gene mapping from Ensembl annotations) and constructed a DESeqDataSet with condition as the design factor. Genes with zero effective length across samples were filtered out prior to analysis. Differential expression testing was conducted in DESeq2 (betaPrior = FALSE) with Benjamini– Hochberg adjustment. Results were further filtered for baseMean > 1 and visualized via volcano plots (geom_point with custom color/size mappings) and baseMean histograms on a log2 scale.

#### Lineage Reconstruction

##### Somatic calling

Sentieon ^32^ TNhaplotyper2 (202308.01) was used in tumor-normal mode to generate somatic call set between all pairs of entities (iPSC lines and PTA/ResolveOme cells). Next, Sentieon TNfilter was run to generate filtered set of TNhaplotyper2 calls. Strelka2 (v2.9.10) was run in tumor-normal mode between all pairs of entities. Intersection of somatic SNVs and InDels tagged with filter = “PASS” by Sentieon TNfilter and Strelka2 internal filters were selected as candidates for downstream analyses.

##### Germline filtering

CNVpytor was used to identify large deletions and loss of heterozygosity (LOH) events in all entities. Copy number events were determined at bin size of 10kbp using combination of read depth and B-allele frequency. Germline calls from NC0 blood bulk sample was used to import germline SNPs. Outcome of CNVpytor was filtered using delta_BAF metric, which represents the change of B-allele frequency from 0.5. Deletion and LOH events with delta_BAF > 0.4 were removed from genomes of entities. CNVpytor was also used to determine regions of allele dropouts in PTA/ResolveOme cells. CNVpytor’s single_cell_allelic_dropout_2 function was run at bin size of 5kbp to generate an output bed file that excludes genomic regions of allele dropouts.

These regions of large deletions and LOHs (in iPSC and PTA/ResolveOme cells) and allele dropouts (in PTA/ResolveOme cells) were excluded from all-to-all pairwise comparisons using All^2^ allele dropout analysis (ADA) mode. Briefly, All2 ADA mode includes somatic mutations in regions of allele dropouts if the VAF is more than 50%. Candidates with (mosaic score)^2^ + (germline score)^2^ > (0.75)^2^ (representing calls beyond the noise region) were selected for further analyses. This strategy resulted in 128,318 candidate set of rarely shared somatic mutations, 22 frequently shared somatic mutations, and still retained 18,576 mutations in likely-germline category.

##### Filtering low frequency amplification artefacts

Next, we determined combined VAF for shared mutations (>= 2 entities) by calculating combined alternate allele supporting reads and combined total depth across all entities in which a mutation is identified. Using exact binomial test, we tested if combined VAF for each shared mutation is significantly lower than expected 50% VAF distribution of heterozygous somatic mutations (this scenario also accommodates somatic calls in regions of allele dropouts as VAF will be much higher than 50%). Mutations with significantly low VAF distribution (p-value <= 0.05) were removed from analyses as these represent low frequency amplification artefacts. This strategy resulted in 52,175 shared mutations, 12 frequently shared mutations, and 14,376 likely-germline mutations.

##### Initial somatic scaffolding based likely-germline filtering

We removed mutations that were identified in all entities but missing in only one, as these likely represent germline mutations that were not called in only one entity by chance (insufficient sampling/unprofiled dropout). Since we previously demonstrated extreme imbalance in NC0’s early lineage contributions, we expanded our focus on candidates from All^2^ with mosaic score < 0.20 (**Fig. S5A**). First, we removed mutations which were observed frequently in population using gnomAD v2.1.1. Only candidates with gnomAD AF < 0.1% and gnomAD filter = “PASS” were retained. This strategy resulted in 51,102 rarely shared mutations, 7 frequently shared mutations, and 524 likely-germline mutations.

Using 51,109 frequently or rarely shared mosaic candidates, we generated an initial tree scaffolding using Jaccard, Dice, Ochiai and Kulczynski2 distances. These distance metrics were selected as they only use shared 1s (shared mutations) to generate corresponding similarity indices between entities, while ignoring shared 0s (shared absences of mutations), with the underlying premise of considering mutations as rare events to be prioritized. We empirically observed that these distances were robust at defining the first two branches in the tree with mutations shared across many cells, even when a few early mutations (that fall into “likely germline” category) are not utilized for the reconstruction. However, they were not robust at constructing later branches when number of shared mutations is smaller and could be comparable to residual false shared calls from amplification. Therefore, using each metric’s computed distance as an initial tree, we query 524 calls in likely-germline category against each tree. Specifically, for each distance-metric based initial scaffold, we query mutations iteratively at each node and identify mutations that fit the shared mutation pattern of the initial scaffold. The requirements for a mutation to be selected are 1) the mutation should be exclusively present on one branch (out-branch threshold = 0), and 2) within a branch the mutation should be seen in at least 80% of its members (in-branch = 80). We select 80% as the in-branch threshold to accommodate for cells with dropouts.

Using this strategy, initial trees generated by all four distance metrics resulted in the same two mutations to be retained from the likely germline category. We used these 2 rescued mutations along with 51,109 somatic mutations to construct the final lineage tree.

##### Lineage reconstruction and comparison

Using final call set of 51,111 somatic mutations, we reconstructed NC0’s lineage distribution with MPBoot ^33^. We observed that with most of the falsely shared mutations removed, the maximum parsimony principle utilized by MPBoot results in the correct topology for early and late branches. Our protocol was able to assign unique mutations to all except two branches of the final lineage tree. The two branches without uniquely characterizing mutations are denoted as ambiguous branches (**Fig. 4G, S6A)**.

In comparing NC0’s de novo reconstructed lineage to Fasching et al. (2021) ^10^, our protocol correctly identified and retained all early mutations from the first division resulting in dominant and recessive lineages (**Fig. S6B**). In the dominant lineage, we further retained all previously identified mutations except ‘n’ (excluded as it’s combined VAF distribution is significantly lower than 50%) and ‘o’ (excluded as gnomAD filter ≠ “PASS”). Additionally, we were able to resolve branching at mutation ‘g’. Notably, we added several internal branches providing higher resolution of middle and late developmental stages in dominant lineage. In the recessive lineage, we identified one additional mutation along with correctly retaining 9/12 previously identified mutations (ε was excluded due to high gnomAD AF; χπ and λ were excluded due to falling within the noise arch).

##### Manual expansion of previous tree

We compared NC0’s *de novo* reconstructed lineage with the manual expansion of known tree constructed previously ^10^ (**Fig. 4G, S6A**). Before expanding the tree, we removed sites with gnomAD AF >= 0.1%. The expansion relied on unfiltered mutation calls from all-to-all pairwise comparisons without allele drop-out mode. We assigned cells to early branches from genotyping early mutations. Reconstruction of later branching was conducted manually based on the number of shared calls. Such expansion suffered from germline contamination and low frequency amplification artefacts resulting in high signal-to-noise ratio of shared mutations between all pairs of entities. We developed the *de novo* reconstruction protocol to address these concerns and provide a comprehensive method to identify early mutations along with accurate resolution at middle and late developmental stages.

## Supplementary Tables

**Table S1. Sample metadata, and somatic variants; related to Figures 1**, **3**, **4**, **5**. T1, Metadata for all samples included in study. T2, SCELLECTOR results for amplification QC. T3, Sequencing metrics, mutation burdens and annotations. T4, number of private and shared mutations identified during lineage reconstruction.

**Table S2. RNA yields and Transcriptome data analysis; related to Figure 2**. T1, RNA yields for standard and modified ResolveOme protocol applied on cells or nuclei. T2, List of differentially expressed genes for nuclei vs cells. T3, GSEA for differentially enriched GO terms and pathways.

## Supplementary Figure

**Fig. S1:**
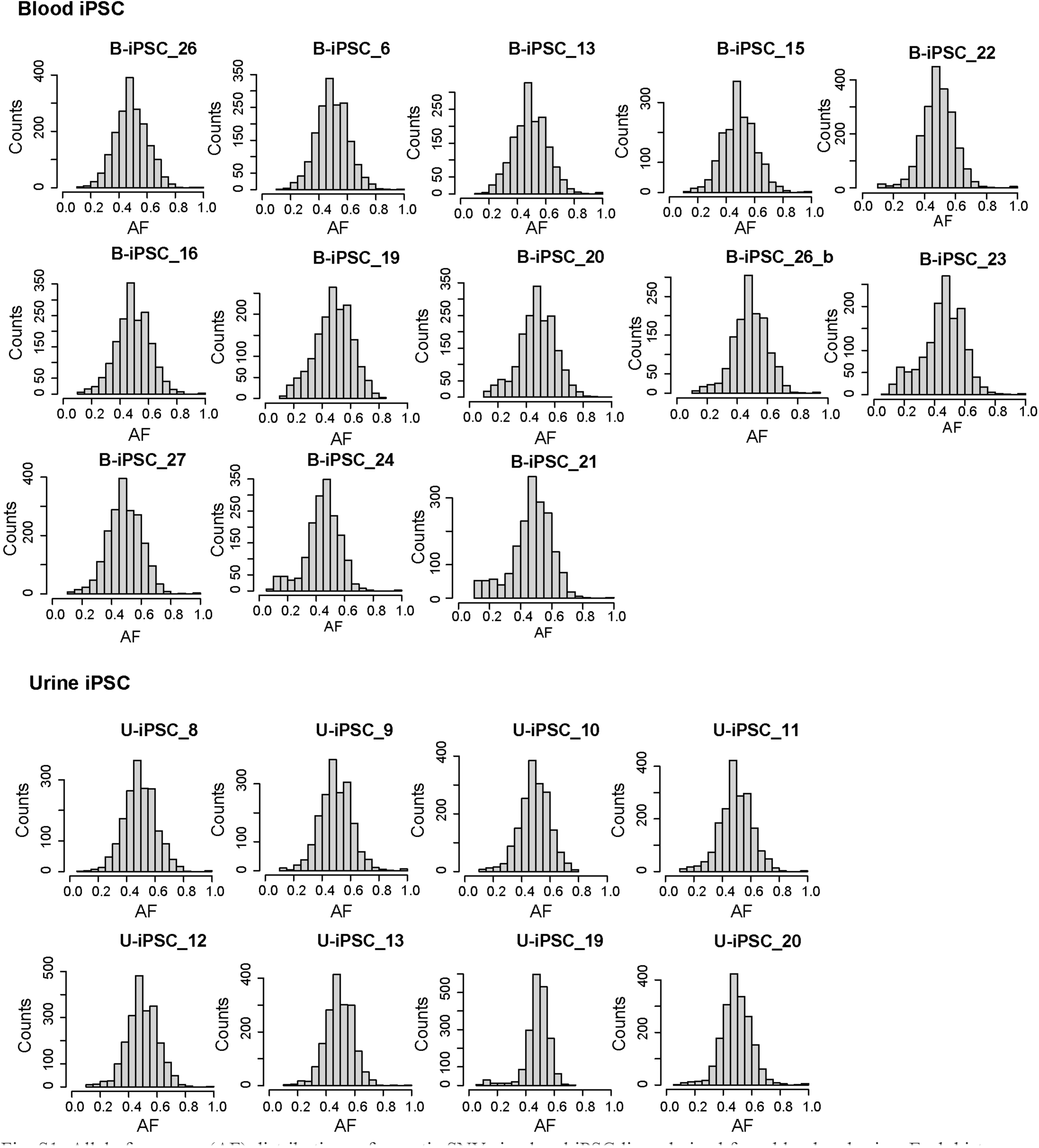
Allele frequency (AF) distributions of somatic SNVs in clonal iPSC lines derived from blood and urine. Each histogram represents the AF profile of somatic SNVs detected in individual iPSC samples from blood and urinary tract.

**Fig. S2:**
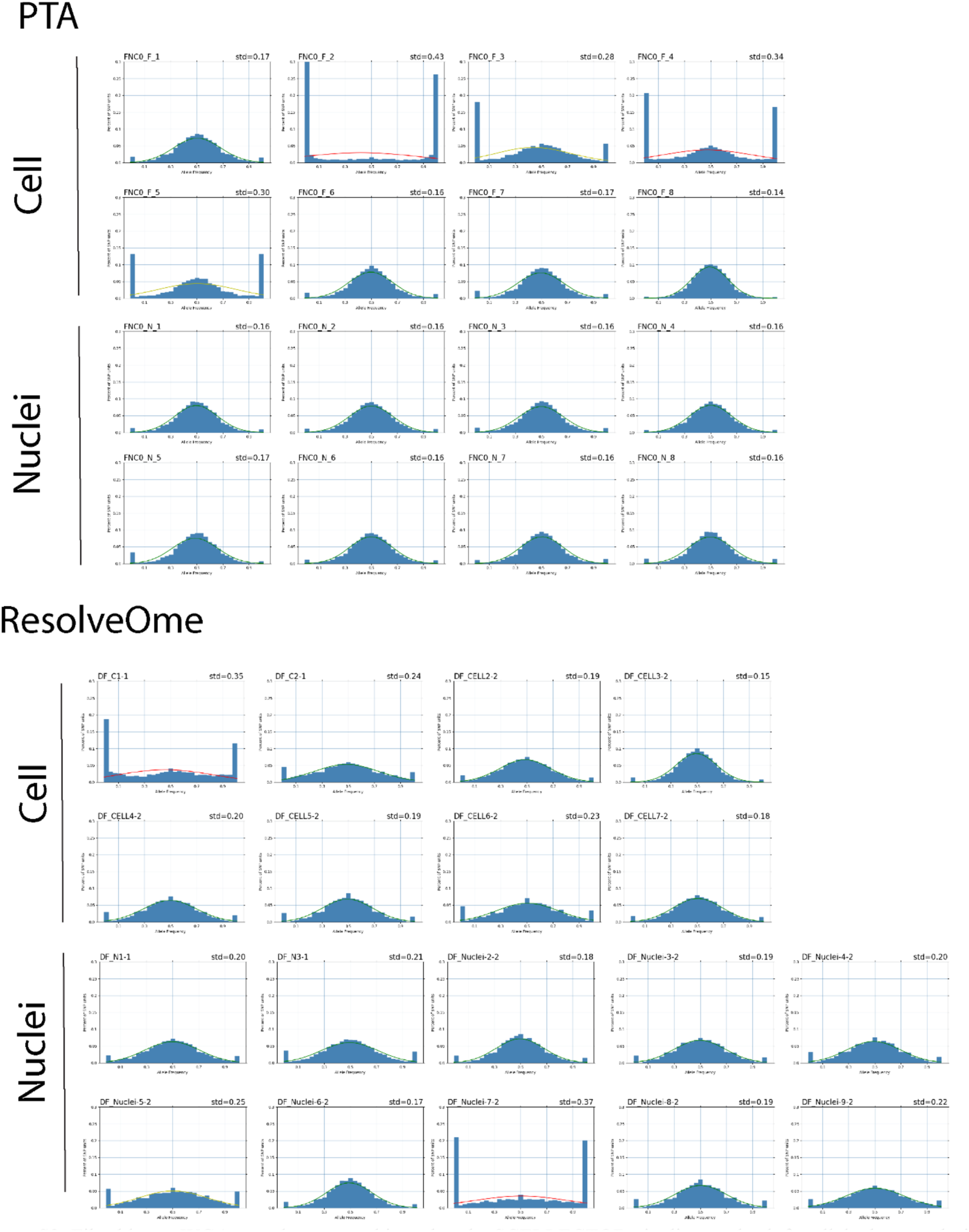
Fibroblast scWGA samples were subjected to the SCELLECTOR pipeline to check for allele dropout during amplification. Each plot represents amplification results for one cell/nuclei. Standard deviation was calculated based on observed allele frequency. Plots with red lines signify failed amplification.

**Fig. S3:**
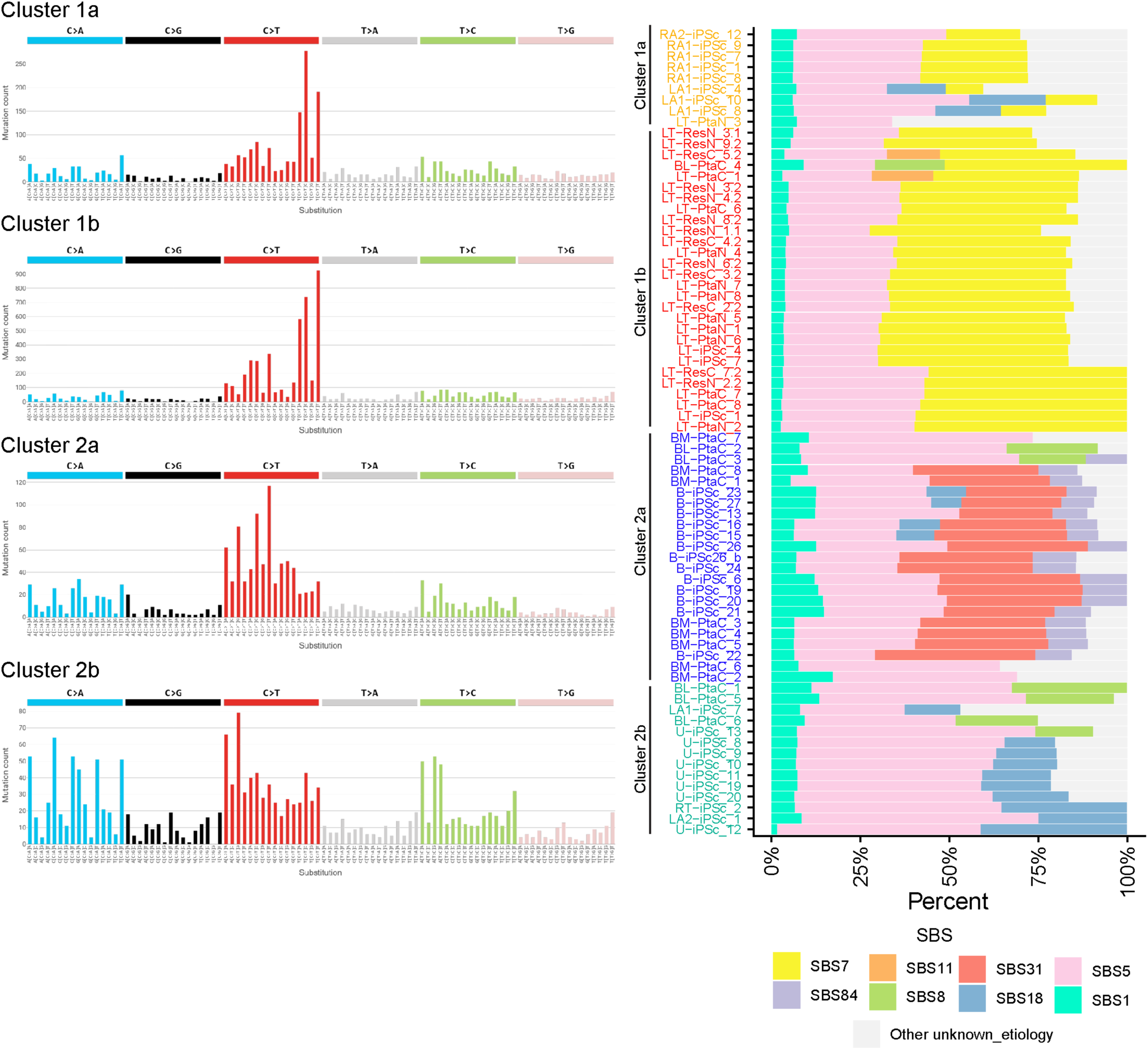
Left panel: Trinucleotide mutation spectra for representative samples from Cluster 1a, 1b, 2a, and 2b. Right panel: Percent stacked bars showing deconvolved contributions of mutational signatures for samples grouped by cluster; colour reflects the fraction of mutations attributed to each signature.

**Fig. S4:**
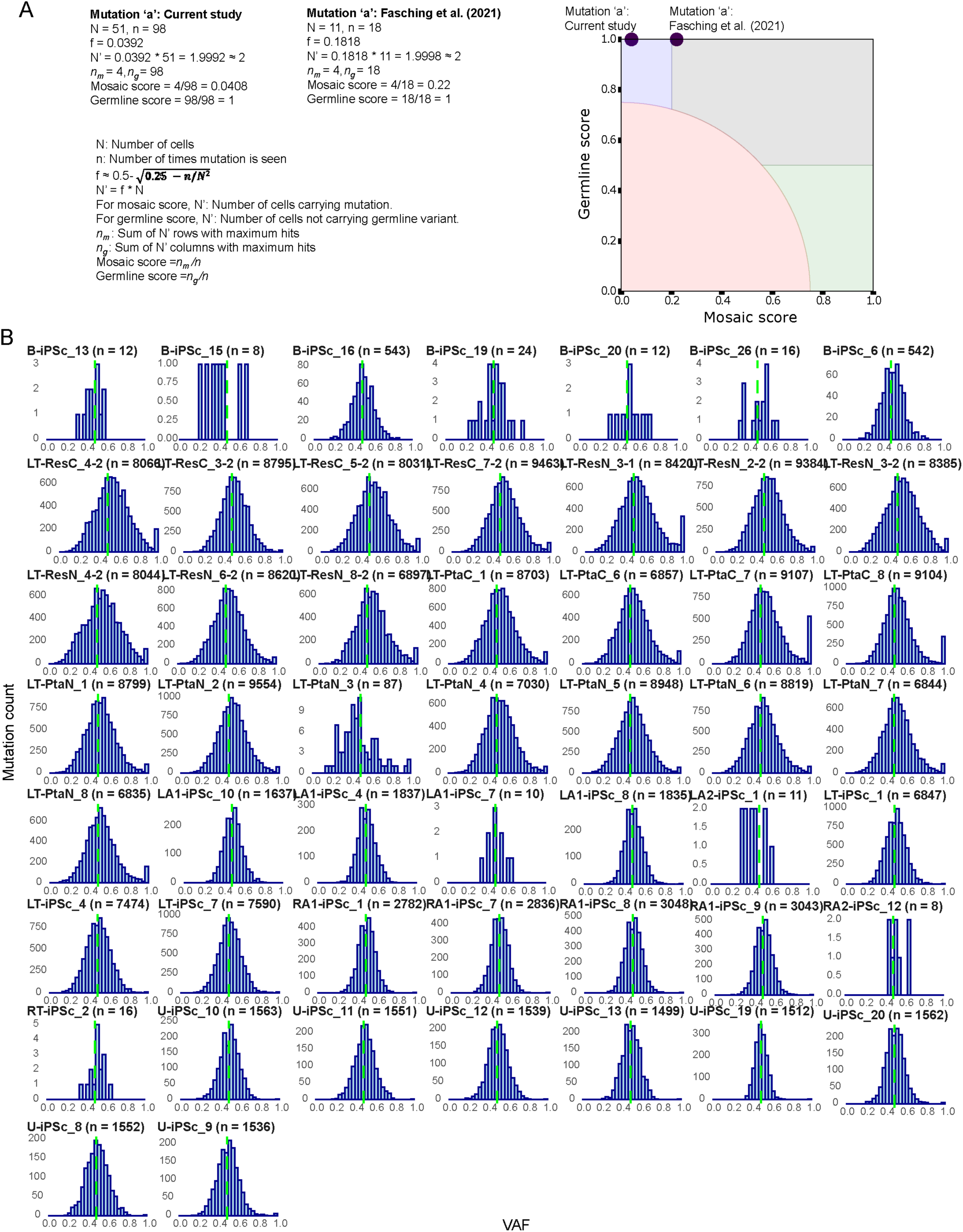
**(A)** Calculation of germline and mosaic scores of mutation ‘a’ (marking dominant lineage in Fig. 4G and Fig. S5A) in Fasching et al. (2021) study placing it in frequently shared region; compared to the current study placing it in the likely germline region. **(B)** Per-entity VAF distribution of final call set of shared mutations used for *de novo* lineage reconstruction centres at 50%. Cells LT-PtaC_7 and LT-PtaC_8 have higher proportion of mutations at VAF = 1 due to shared loss of chromosome X (Fig. 5B). Entities with few shared mutations indicate early divergence.

**Fig. S5:**
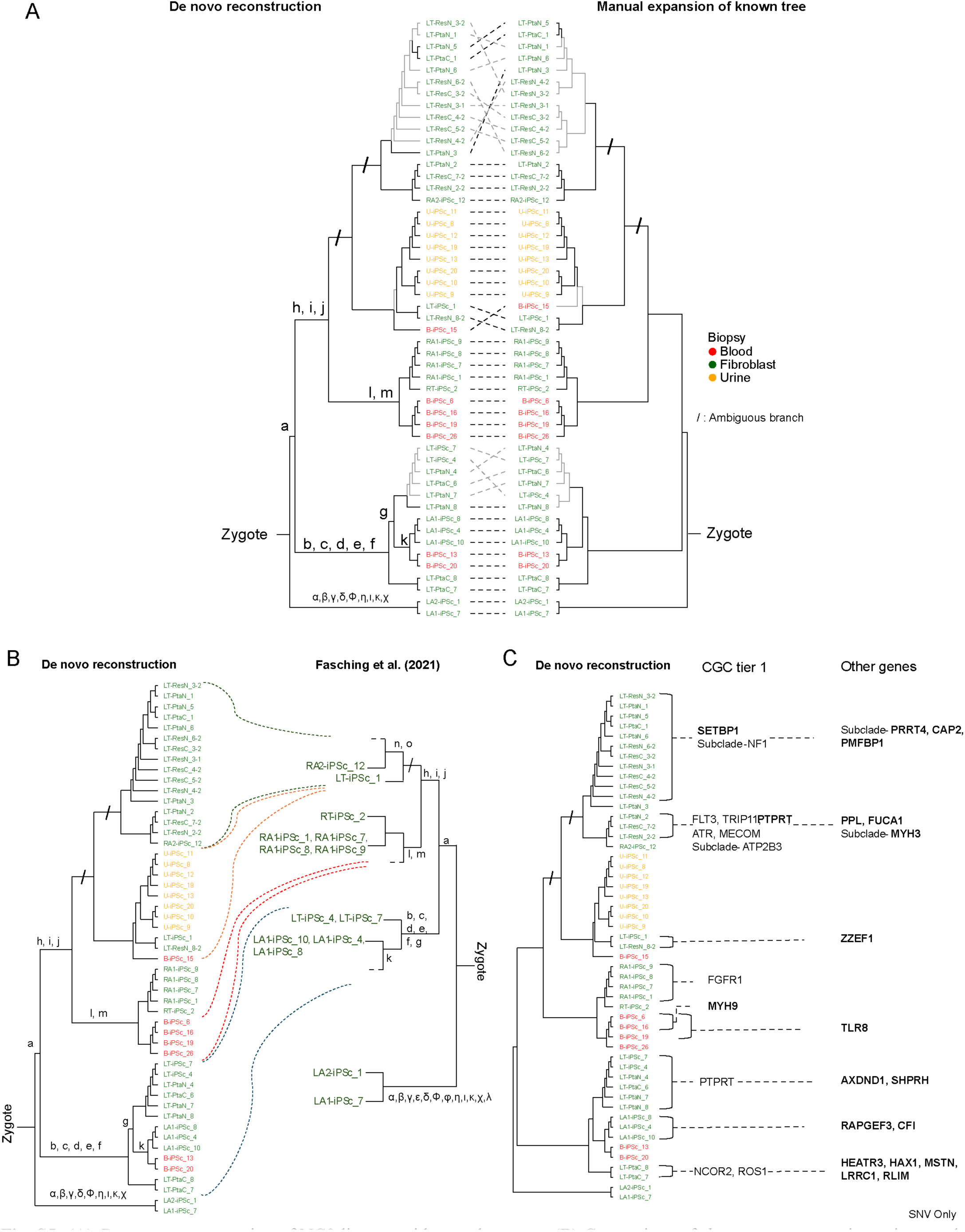
**(A)** *De novo* reconstruction of NC0 lineage with sample names. **(B)** Comparison of *de novo* reconstruction using newly developed method against lineage analysis in Fasching et al. (2021) reveals accurate identification of early mutations and higher resolution of branching within dominant lineage. **(C)** Functional annotation of shared missense and truncating SNVs identified within coding regions of Cancer Gene Census (CGC) Tier 1 set and other genes, mapped to *de novo* reconstructed branches. Genes in bold indicate presence of truncating SNVs, while genes not in bold indicate presence of missense SNVs. Genes not in CGC-Tier 1 set were prioritized for visualization if they harbor truncating mutations.

**Fig. S6:**
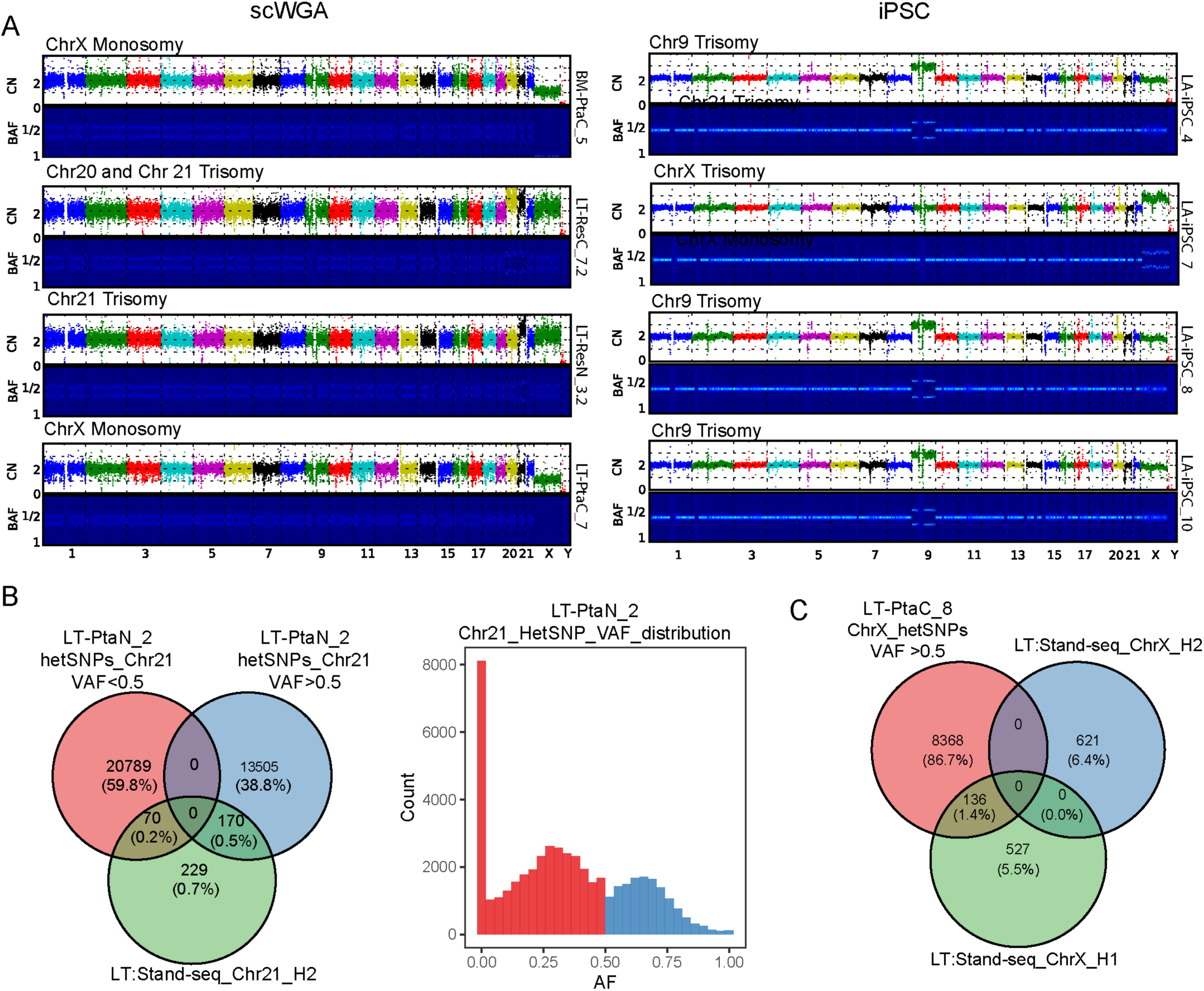
A) Read-depth plots from scWGA-cells and iPSC lines highlighting copy-number variation: chr21 trisomy, chr20 trisomy chrX monosomy, chr15 monosomy, chr9 trisomy and chrX trisomy. B) Venn diagram representing overlap between Strand-seq phased heterozygous SNPs assigned to haplotype H2 and variants observed in the PTA sample LT-PtaN_2 (chr21 trisomy). Scatter of hetSNPs is coloured by PTA alternate-allele fraction (VAF ≤ 0.5, red; VAF > 0.5, blue). Inset histogram shows the VAF distribution of chr21 hetSNPs in LT-PtaN_2, illustrating the shift of H2-matching sites toward higher VAF consistent with duplication of H2. C) Intersection of Strand-seq phased X-chromosome hetSNPs (H1 and H2) with variants detected in LT-PtaC_8 (chrX monosomy). Venn diagram highlights exclusive overlap with H1-phased SNPs, consistent with loss of haplotype H2 in the monosomic cell.

**Fig. S7:**
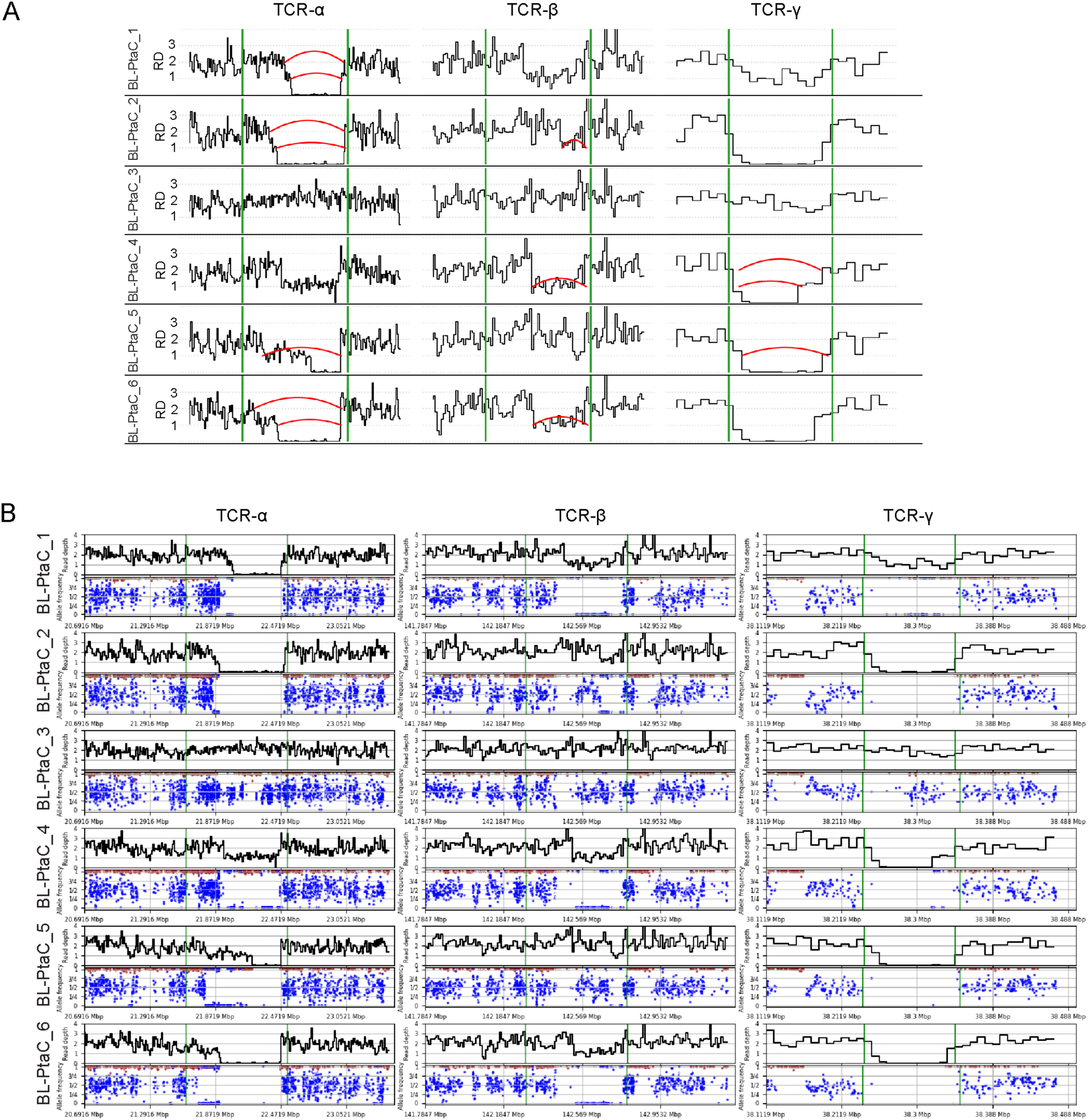
(A) Locus-level read-depth and junction signals for TCR-α, TCR-β, and TCR-γ across lymphocyte samples, red arcs highlighting putative V(D)J rearrangements and green vertical lines marking canonical segment boundaries used to orient calls. (B) Locus read-depth signal and supporting allele frequency of heterozygous SNPs plotted below (blue dots), enabling visual confirmation of rearrangement breakpoints and read support. Change in copy number, and split-read clusters distinguishes rearranged lymphocytes (e.g. BL-PtaC_2) from non-rearranged samples (flat coverage; BL-PtaC_3), permitting assignment of cell identity (T-cell versus NK cell).

**Fig. S8:**
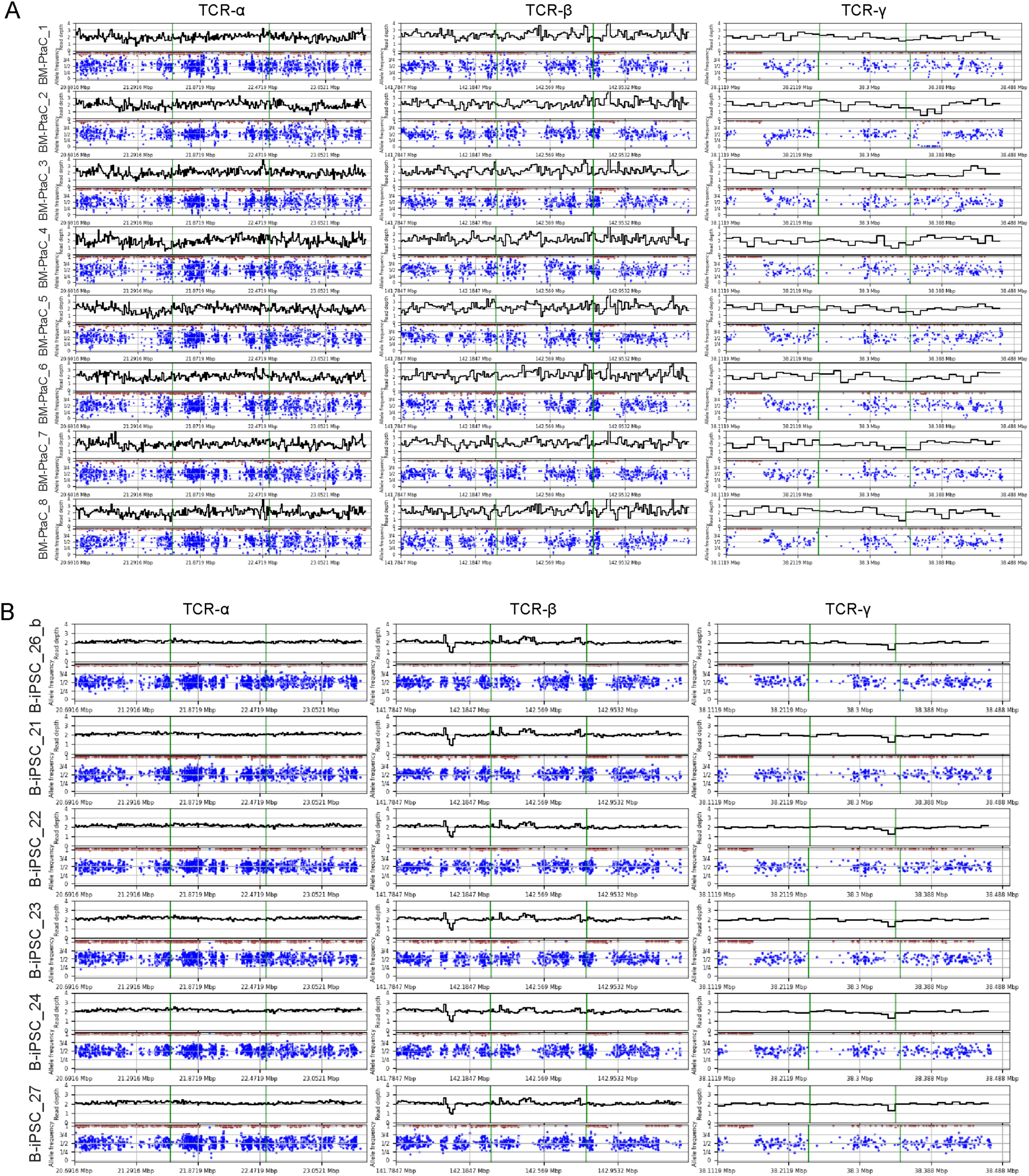
Locus read-depth (top) and supporting allele frequency of heterozygous SNPs (bottom) enabling visual confirmation of lack of rearrangement (flat coverage) at the TCR locus for blood monocytes (A) amplified by PTA and blood-derived iPSCs (B).

